# Revealing the platelet transcriptome using Siphon-seq: a sensitive, purity-aware method for RNA-sequencing

**DOI:** 10.64898/2026.11.16.767104

**Authors:** Brittany A. Moser, Catharina Conrad, Qin Zhu, S. Farshid Moussavi-Harami, Catherine A. Taylor, Longhui Qiu, Carolyn S. Calfee, Zev J. Gartner, Mark R. Looney

**Affiliations:** Department of Pharmaceutical Chemistry, UCSF; San Francisco, California, USA; Department of Pediatrics, UCSF; San Francisco, California, USA; Department of Medicine, UCSF; San Francisco, California, USA; Chan Zuckerberg Biohub; San Francisco, California, USA

## Abstract

Platelet transcriptomics is limited by the exceptionally low RNA content of platelets and by disproportionate contributions from contaminating cells to bulk RNA-sequencing libraries. We developed Siphon-seq, an approach that deliberately overloads independently barcoded single-cell partitions to increase aggregate platelet RNA input, then computationally identifies and removes partitions containing contaminating RNA after sequencing. Applied to 34 healthy human donors, Siphon-seq increased platelet-marker representation, depleted leukocyte and erythrocyte transcripts, and removed contamination-associated inter-donor transcriptional structure. We defined a 3,436-gene core platelet transcriptome that was strongly related to megakaryocyte expression. However, megakaryocyte transcription did not fully explain the platelet transcriptome, which had specific pathways systematically enriched or depleted that were obscured by leukocyte contamination. These findings were confirmed in an independent cohort and were associated with protein detection in two platelet proteome resources. In a pilot human sepsis cohort, Siphon-seq reduced transcripts associated with leukocytes to reveal increased platelet-associated pathway signals. These contaminating immune pathways were also enriched in samples analyzed by bulk RNA-seq from an independent sepsis cohort, most strongly in the stratum with the highest estimated leukocyte burden. Combined, these results reveal that RNA-level contamination can alter both transcriptome structure and biological inference, and that Siphon-seq bridges the sensitivity-purity tradeoff that has limited platelet RNA sequencing, enabling sensitive, purity-aware studies of platelet biology in health and disease.

## Introduction

Platelets are anucleate blood cells essential for hemostasis, thrombosis, and immune responses.^1–3^ Despite lacking a nucleus, they retain a functional transcriptome of mRNAs and non-coding RNAs inherited from megakaryocyte progenitors that are translated and spliced in response to physiological stress and disease.^4–6^ This RNA pool is of interest both as a probe of platelet biology and as a circulating biomarker source in cardiovascular disease, cancer, and infection.

Profiling platelet RNA accurately and at scale remains technically challenging, and there is no clear consensus on how the platelet transcriptome changes with aging or across disease contexts.^6–9^ The most common approach, bulk RNA sequencing of platelets isolated from platelet-rich plasma, depletes most erythrocytes and leukocytes by sequential centrifugation, but residual contaminating cells and debris may remain in the final pellet.^10–12^ A single platelet contains 1,000- to 12,500-fold less RNA than a leukocyte, so even trace contamination distorts platelet libraries. In platelet microarray data, low-level leukocyte contamination has been shown to alter over 8,000 transcripts, artificially enriching immune and cytokine signaling while suppressing genuine platelet signal.^12,13^ This problem compounds in disease, since platelet-leukocyte aggregates and leukocyte-derived microparticles increase with inflammatory burden in sepsis and malignancy, making contamination correlate directly with the biological comparison of interest.^14–17^

Single-cell RNA-seq could resolve RNA contamination by isolating platelets and other cells in separate droplets, but standard droplet-based platforms such as 10x Chromium are poorly suited for platelets.^18^ Platelet RNA content (∼2.2 fg) falls two to three orders of magnitude below the input typically required for robust library generation. At such low template levels, adapter-dimer products dominate over genuine cDNA, a failure we confirmed across five commercial chemistries at maximum loading (**Supplemental Table 1**).^7,19^ Platelets also aggregate upon activation, and standard doublet-detection methods, which assume similar RNA content across cells, cannot reliably identify platelet aggregates because platelet RNA content varies widely with maturity. ^18,20–25^ Transcriptionally distinct platelet subpopulations have been proposed, but low per-platelet RNA limits transcript detection, and without orthogonal validation, aggregates or multiplets cannot be excluded as contributors to the observed heterogeneity. ^17, 18,26–28^

Rather than profile single platelets, we used single-cell technology to generate bulk platelet transcriptomes. We hypothesized that we could obtain clean bulk platelet transcriptomes by inverting standard single-cell practice: deliberately overloading GEMs in the 10x Chromium workflow so each captures many platelets rather than one, raising aggregate RNA input above the biochemical failure threshold while preserving independent per-GEM barcoding. Barcoding shifts the contamination problem to one resolved digitally, by identifying and excluding partitions bearing high-confidence non-platelet marker expression, including leukocyte-platelet aggregates that evade physical removal, yielding a digitally purified pseudobulk platelet transcriptome without requiring single-platelet RNA yields. We term this approach **Siphon-seq**: platelet-rich plasma is conventionally prepared by centrifugation, separating the platelet-rich supernatant from a pelleted fraction enriched for leukocytes and erythrocytes. Siphon-seq performs an analogous separation digitally, computationally “siphoning” platelet-enriched partitions from those containing RNA from contaminating cell types.

Here, we begin by providing evidence that many published platelet RNA-seq datasets contain variable contributions from contaminating cell types that affect data interpretation. We then apply Siphon-seq to 34 healthy human donors, defining a core human platelet transcriptome that closely tracks megakaryocyte expression, but has systematic deviations normally hidden by leukocyte contamination. We then extend Siphon-seq to fixed clinical samples in a pilot sepsis cohort. There, we demonstrate that contamination alters pathway-level inference, both introducing leukocyte-associated signals and obscuring platelet-associated responses.

## Methods

### Study participants

The healthy cohort comprised 34 adults (median age 36.5 years, interquartile range 31–47, range 23–59; 17 male and 17 female; self-reported race/ethnicity: 17 White, 11 Asian, 4 Middle Eastern/North African, 2 Hispanic/Latino) (**Supplemental Table 2**). The fixed-RNA pilot comprised 8 participants: 4 patients with sepsis and 4 controls, of whom 2 were critically ill patients without sepsis and 2 were healthy volunteers (**Supplemental Table 3**). Patients with sepsis and critically ill controls were enrolled in the Early Assessment of Renal and Lung Injury (EARLI) cohort under UCSF IRB protocol #10-02852, which granted a waiver of initial consent for blood sampling.^29,30^ Consent was subsequently obtained from patients or their surrogates where possible; otherwise, patients were included under an IRB-approved waiver of consent. Healthy volunteers gave written informed consent under UCSF IRB protocol #24-41921. The study was conducted in accordance with the Declaration of Helsinki.

### Platelet isolation

Whole blood was collected into EDTA vacutainer tubes. Platelet-rich plasma (PRP) was prepared by centrifugation at 300 × g for 10 min with acceleration and deceleration set to zero, and supplemented with prostacyclin (PGI₂, 0.1 μg/mL) and apyrase (0.25 U/mL) to limit activation.^31^ Platelets were pelleted at 1000 × g for 10 min and resuspended in HEPES-buffered Buffer A (pH 6.7–6.8) containing the same inhibitors, then passed through a 20-μm filter to remove residual nucleated cells and aggregates. Platelet concentration was determined by flow cytometry using CD41 staining and CountBright Plus counting beads.

### GEM overloading

Platelets were loaded far above the standard 10x Chromium target, at 5 × 10⁶ per channel for Next GEM Single Cell 3′ v3.1 and 1 × 10⁷ for GEM-X Single Cell 3′ v4, so that each GEM captured many platelets while retaining an independent partition barcode.

### Siphon-seq classification and pseudobulk

Sequencing reads were processed with Cell Ranger 8.0.1 against the GRCh38-2024-A human reference (mm10 for mouse) at default settings. Partitions were called from the inflection point of the counts-per-partition curve. Counts were normalized and embedded in Scanpy (nearest-neighbor graph, UMAP, Leiden clustering), and leukocyte, erythrocyte and platelet populations were annotated.^32–34^ The top 500 differentially expressed genes per population were used as features to train a CONCORD classifier, which assigned each partition a type together with leukocyte (WBC_prob) and erythrocyte (RBC_prob) probabilities.^35^ Platelet partitions above a contaminant-probability threshold were relabeled high-background, shown as a separate category, and excluded from the Siphon-seq pseudobulk. Purified pseudobulk summed raw UMI counts across retained platelet partitions within each donor; pre-purification pseudobulk summed all partitions from that donor, giving a matched within-donor contrast. Thresholds and clustering rules are given in the Supplemental Methods.

### Core transcriptome and comparisons

Donor pseudobulk counts were TMM-normalized (edgeR).^36,37^ Genes averaging below 2 CPM were treated as below the detection floor, and genes exceeding 2 CPM in ≥90% of donors defined the 3,436-gene core transcriptome. Whether a gene reached the core was modelled by logistic regression on megakaryocyte expression (log₂(FPKM + 1); platform-matched BLUEPRINT RNA-seq of CD34−CD41+CD42+ cord-blood megakaryocytes, n = 6), giving an expected core count for each gene set against which the observed count was compared.^38,39^ An extended model adding mRNA half-life and nuclear:cytoplasmic ratio is reported in the Supplemental Methods. Cell-type RNA composition of published bulk platelet datasets was estimated with MuSiC 1.0.0 (music_prop, default arguments) against a single-cell reference and expressed as RNA share.^40^

### Differential expression and pathway analysis

Differential expression between sepsis and control was computed separately within the pre- purification and Siphon-seq preparations (pyDESeq2), and genes ranked by the Wald statistic were tested by preranked gene set enrichment analysis against GO Biological Process 2023 (gseapy).^41,42^ The effect of purification on each pathway was summarized as the change in normalized enrichment score between preparations, ΔNES = NES_Siphon-seq − NES_pre-purification. Gene filters, contrasts, thresholds and permutation settings are given in the Supplemental Methods and software and version numbers are listed in **Supplemental Table 4**.

### H2Data sharing statement

Sequencing data generated in this study will be available upon manuscript acceptance. Analysis code will be available on Github upon manuscript acceptance. Publicly available data reanalysed here: 14 bulk platelet RNA-seq projects from PlateletBase (BioProject, ENA and GEO accessions in Supplemental Table 5); GSE163668 and GSE184916 (deconvolution reference); GSE252275 (independent sepsis cohort); GSE85864 and GSE94384 (platelet translation reference); and BLUEPRINT RNA-seq (European Nucleotide Archive ERP001466 and ERP001664).

## Results

### Siphon-seq removes contamination-driven transcriptional structure from bulk platelet RNA-seq

We first investigated whether published bulk RNA-seq data of human platelet transcriptomes bore hallmarks of leukocyte or erythrocyte contamination using MuSiC transcriptome deconvolution. Because no validated platelet scRNA-seq reference exists, we built a six cell-type reference from a whole-blood dataset, adding erythroid profiles from a dataset in which red blood cells were captured directly, since the whole blood dataset did not recover mature erythrocytes. We found that cell-type composition varied widely across bulk platelet RNA-seq samples, even among preparations with high median platelet RNA (**Fig. 1A, Supplemental Figure 1, Supplemental Data 1**).

**Figure 1.**
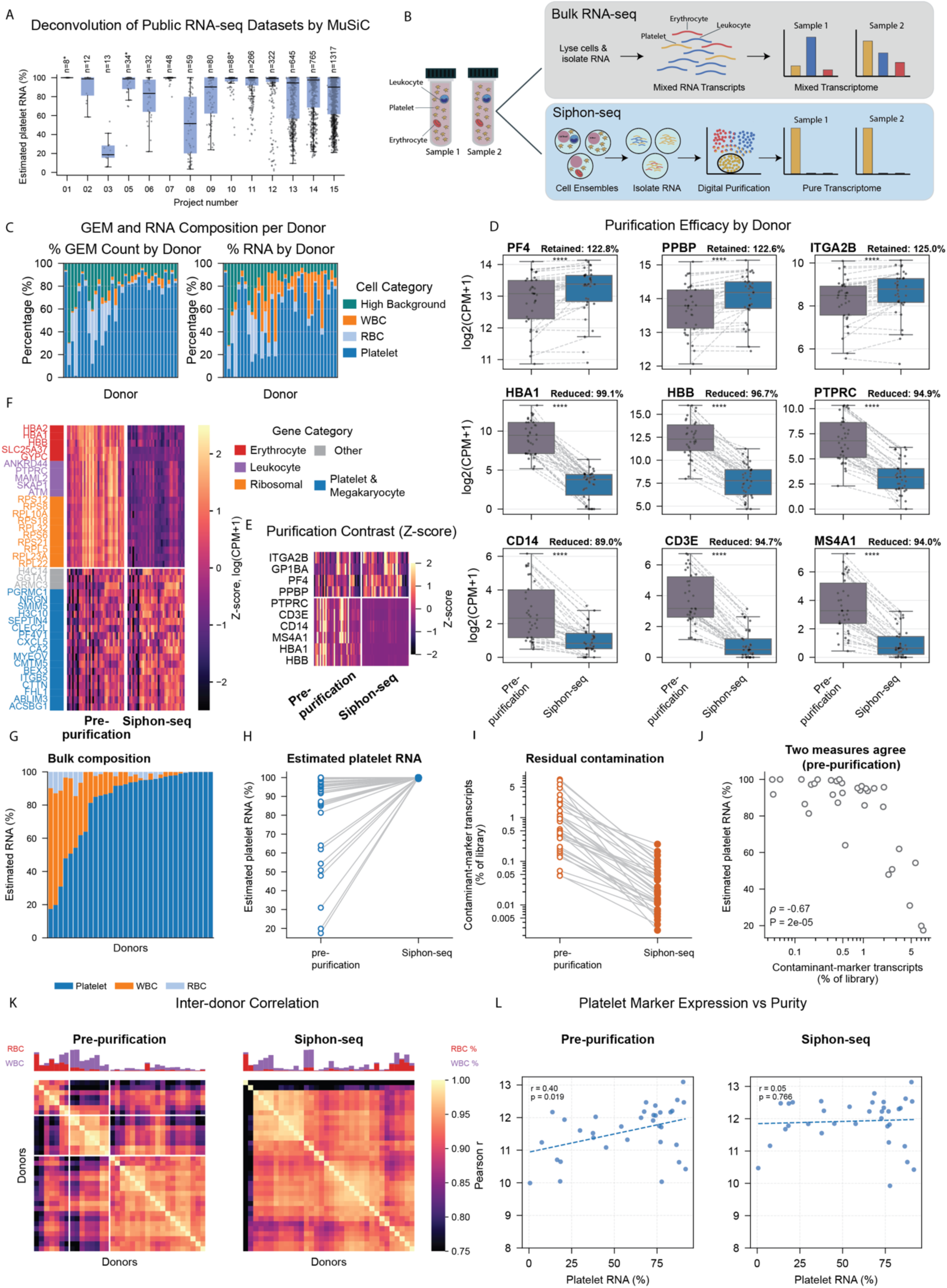
Siphon-seq removes contamination-driven transcriptional structure from bulk platelet RNA-seq. **(A)** MuSiC-estimated platelet RNA fraction for 3,689 samples across 14 public PlateletBase bulk RNA-seq projects (accessions in Supplemental Table 5). Sample size (n) given above each box; asterisks (*) indicate CD45-depleted platelet isolation. **(B)** Schematic comparing conventional bulk RNA-seq, where RNA from all cell types is pooled before sequencing, with Siphon-seq, which uses single-cell ensembles and computational classification of partitions to isolate platelet-specific transcriptomes from mixed preparations. **(C)** Per-donor cell type composition (left, by cell count) and RNA composition (right, by total RNA) in pre-purification preparations across our 34-donor cohort. Colors indicate platelet, erythrocyte (RBC), leukocyte (WBC), and high-background (platelet GEMs with residual contaminant signal). **(D)** Matched-pair comparison of platelet markers (PF4, PPBP, ITGA2B) and contaminating markers (HBA1, HBB, PTPRC, CD14, CD3E, MS4A1) in pre-purification versus Siphon-seq preparations from the same donors. log2(CPM+1); dashed lines connect matched pairs. Wilcoxon signed-rank test, **** p < 0.0001; percentages show mean retention (platelet markers) or reduction (contamination markers) after purification. **(E)** Z-score heatmap of the same canonical markers across pre-purification and Siphon-seq preparations. **(F)** Genome-wide differential expression between pre-purification and Siphon-seq preparations from the same 34 donors, genes annotated by lineage category (Supplemental Data 11). **(G)** MuSiC-estimated cell type fractions (platelet, WBC, RBC) for our 34-donor dataset prior to Siphon-seq. **(H)** MuSiC-estimated platelet fraction per donor, pre-purification versus Siphon-seq, confirming near-complete purity by an independent deconvolution method. **(I)** Combined CPM of the six contaminating markers from (D), pre-purification versus Siphon-seq. **(J)** Spearman correlation between MuSiC-estimated purity and contaminant-marker abundance across donors before purification, cross-validating the two contamination measures. **(K)** Inter-donor Pearson correlation matrices (log2 CPM, all expressed genes), pre-purification (left) and Siphon-seq (right), each matrix ordered by its own hierarchical clustering. Annotation bars show per-donor RBC/WBC contamination; white lines demarcate bulk donor clusters by predominant contamination type. Color scale truncated at r = 0.75. **(L)** Per-donor platelet purity versus mean platelet marker expression (PF4, PPBP, ITGA2B), pre-purification (left) and Siphon-seq (right). Dashed line: linear regression; Pearson r and p-value shown.

We applied Siphon-seq to platelets from 34 healthy donors to generate a high-purity consensus platelet transcriptome. Platelets were enriched from whole blood by centrifugation. GEMs were then loaded more than two orders of magnitude above the standard single-cell recommendation, so that each partition captured an ensemble of many platelets. Partitions containing non-platelet RNA were then identified and excluded (**Fig. 1B, Supplemental Figure 2**). The fraction of GEMs classified as pure platelet ensembles ranged from 1.4% to 93.9% across donors (median ∼80%). Twenty-four of 34 donors exceeded 70%, and 9 fell below 50%, mainly due to erythrocyte contamination (**Fig. 1C, Supplemental Data 2**). GEM counts underestimated contamination at the RNA level. Leukocyte-containing partitions contributed far more RNA than their numbers predicted, so even a few leukocytes substantially reduced platelet RNA purity (**Supplemental Figure 3**).

For each donor, we compared the pre-purification library (all GEM partitions) with the same library after contaminant partitions were removed (Siphon-seq). Siphon-seq significantly increased the canonical platelet markers PF4, PPBP, and ITGA2B and reduced erythrocyte and leukocyte markers (all P < .0001, paired Wilcoxon; **Fig. 1D–E**). Genes enriched before purification were dominated by ribosomal, leukocyte and erythroid transcripts, whereas Siphon-seq revealed obscured platelet-enriched genes including CMTM5, PF4V1 and CTTN (**Fig. 1F, Supplemental Data 11**). Deconvolution with MuSiC independently confirmed this shift. The estimated platelet RNA fraction varied widely across pre-purification libraries (17.4–100%; **Fig. 1G**) and rose after Siphon-seq (median 93.7% to 100%; higher in 29 of 33 donors, excluding one donor with too few platelet counts after purification, P = 2.1 × 10⁻⁶; **Fig. 1H–J**). Estimated platelet RNA stayed just below 100% in some libraries, likely reflecting small amounts of ambient RNA. The four donors without an increase showed a ceiling effect (99.9–100% before purification). Inter-donor correlations were high in both datasets (median r = 0.91 pre- purification, 0.93 Siphon-seq), but pre-purification libraries clustered donors by contamination type rather than by platelet biology, a structure abolished after Siphon-seq (**Fig. 1K**). Platelet purity correlated with platelet-marker expression before purification but not after (**Fig. 1L**), removing a systematic confounder of cross-donor comparisons. Siphon-seq performed similarly on fresh platelets from a murine sepsis model (**Supplemental Figure 4**).

### Siphon-seq reveals selective departures of the platelet transcriptome from megakaryocyte expression

We next defined a core platelet transcriptome as genes with mean expression ≥2 CPM that exceeded 2 CPM in ≥90% of donors; 3,436 of 38,606 quantified genes met both criteria (**Figure 2A, Supplemental Data 3**). Genes detected above the detection threshold but below the core cutoff were considered contextual, reflecting biological variation; most sex-linked genes fell in this tier: seven Y-linked genes were detected only in male donors, whereas XIST, present at low levels in female donors (mean 0.88 CPM across donors), fell below the detection threshold (**Supplemental Figure 5**). With a clean core transcriptome in hand, we next asked whether its membership reflects selective retention within platelets or simply stochastic sampling from megakaryocyte abundance. Because platelets inherit their RNA from megakaryocytes, we took megakaryocyte expression (BLUEPRINT) as that baseline. Megakaryocyte expression strongly predicted core membership: the core fraction rose from 0.7% in the lowest megakaryocyte-expression bin to 85% in the highest (**Figure 2B, Supplemental Figure 6**). However, it explained only part of core membership (McFadden pseudo-R² = 0.208).

**Figure 2.**
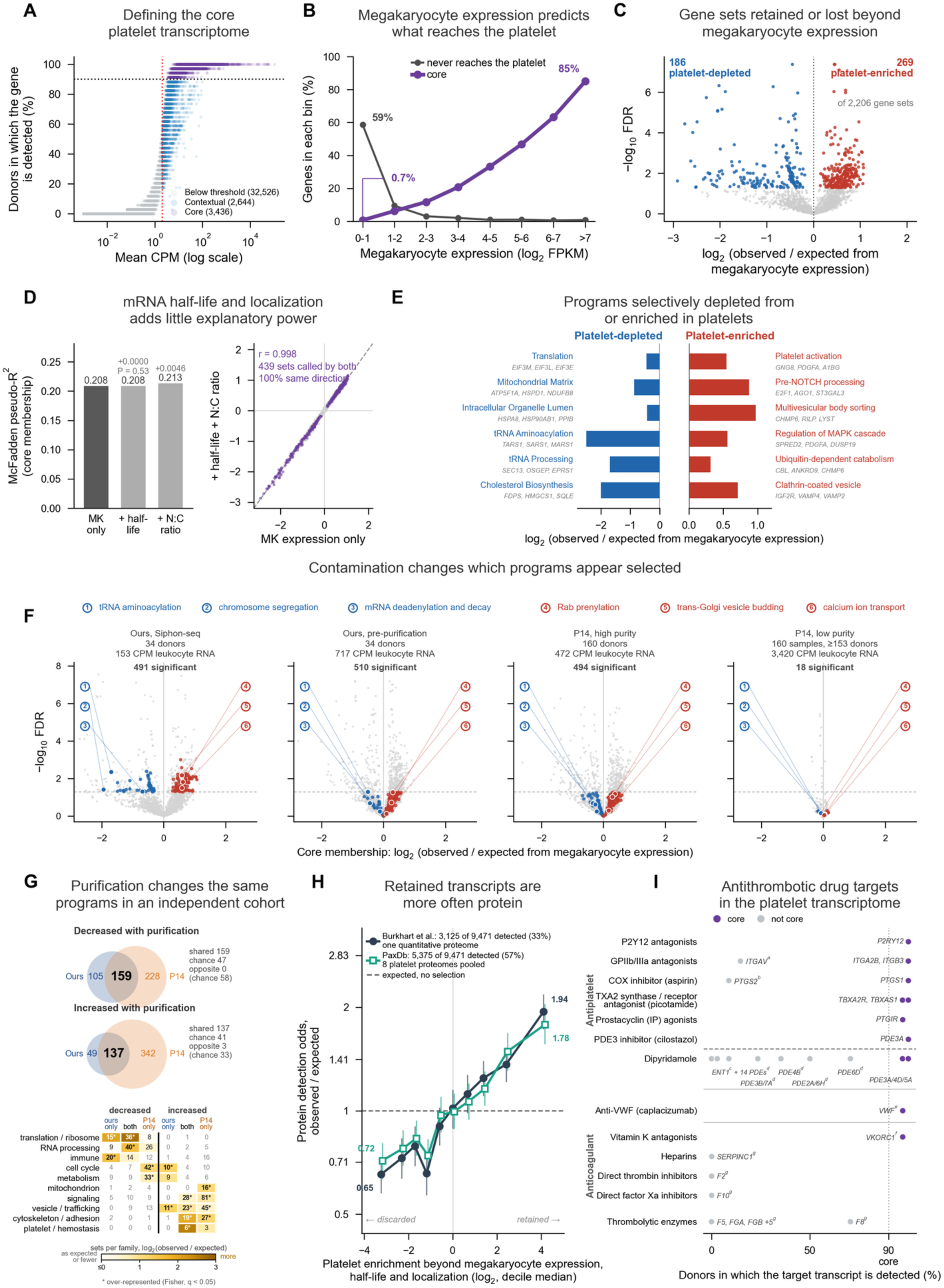
The platelet core transcriptome departs systematically from megakaryocyte expression. **(A)** Mean TMM-CPM across 34 donors versus the percentage of donors in which each gene exceeds 2 CPM. Dotted lines indicate the core criteria (mean ≥2 CPM; detected in ≥90% of donors). Purple, core (3,436); blue, contextual (2,644); gray, below detection (32,526). **(B)** Percentage of genes reaching the core (purple) or never detected in platelets (gray), stratified by megakaryocyte expression (BLUEPRINT, log₂(FPKM + 1)). **(C)** Observed versus expected core genes for 2,206 GO and Reactome sets (≥20 genes); expected values were estimated from megakaryocyte expression by logistic regression. Colors indicate platelet-depleted (blue) and platelet-enriched (red) sets at BH FDR <0.05. **(D)** Left, McFadden pseudo-R² for core membership modeled by megakaryocyte expression alone and after addition of mRNA half-life and nuclear:cytoplasmic (N:C) ratio. Right, per-set log2(observed/expected) under the megakaryocyte-only versus full model; Pearson *r* shown. **(E)** Six most platelet-depleted (left) and platelet-enriched (right) sets from (C) by |z|; gray symbols indicate the three genes contributing most to observed − expected. **(F)** Analysis in (C) repeated across four datasets spanning 153–3,420 CPM leukocyte-marker RNA: Siphon-seq, the same 34 donors before purification, and high- and low-purity halves of PlateletBase P14 (PRJNA659491). Colored points indicate 149 sets significant in purified data but in none of three moderately contaminated datasets (pre-purification and the high-purity halves of P14 and of PlateletBase P13, PRJNA353588; Supplemental Figure 7); numbered points identify six representative programs. **(G)** Gene-set changes with purification in Siphon-seq and P14. Venn diagrams show sets decreased (top) or increased (bottom) with purification at FDR <0.05 and |change| ≥0.25 log2; overlap was tested by Fisher’s exact test. Heatmap shows program-family enrichment within each Venn region; asterisks indicate BH *q* <0.05. **(H)** Observed/expected odds of detectable platelet protein across deciles of platelet transcript enrichment beyond megakaryocyte expression, mRNA half-life, and N:C ratio. Points denote decile medians; bars, 95% bootstrap CIs; dashed line, observed = expected. Protein detection was assessed using Burkhart et al. (33% detected) and PaxDb (57%). **(I)** Human targets of approved antithrombotic drugs (ChEMBL; WHO ATC B01A), plotted by percentage of donors with detectable target transcript. Purple, core; gray, noncore; dotted line, 90% core threshold. Lettered targets primarily transcribed or acting outside platelets are (a) ITGAV, (b) COX-2, (c) ENT1, (d) tissue-specific phosphodiesterase isoforms, (e) VWF, (f) VKORC1, and (g) plasma/endothelial proteins. Drugs, classes, and targets are listed in Supplemental Data 7.

For each GO and Reactome gene set, we compared observed core genes with the number expected from megakaryocyte expression. Of 2,206 sets, 269 contained more core genes than expected and 186 fewer (**F**i**gure 2C****, Supplemental Data 4**), a systematic, bidirectional departure. We next asked whether the departures simply reflected preferential retention of stable or cytoplasmically localized transcripts. They were not explained by mRNA half-life or nuclear-to-cytoplasmic ratio (**Figure 2D**). Platelets were depleted of transcripts for cytoplasmic translation, tRNA aminoacylation, the mitochondrial matrix, and cholesterol biosynthesis, and enriched for platelet activation and aggregation, vesicle sorting, MAPK signaling, and ubiquitin-dependent protein catabolism (**Figure 2E**).

Importantly, contaminating transcripts masked deviations from the megakaryocyte transcriptome. In all pre-purification libraries and in the high-purity half of a public platelet cohort, similar numbers of gene sets reached significance as with Siphon-seq, but the median |log₂(observed/expected)| fell from 0.57 to 0.21–0.23, and to 0.03 in the low-purity half (**Figure 2F, Supplemental Figure 7, Supplemental Figure 8**). In silico mixing of Siphon-seq and pre- purification profiles reproduced this decline, indicating that purity itself drove the attenuation (**Supplemental Figure 9**). The effects of purification were concordant across cohorts (r = 0.87; **Figure 2G, Supplemental Data 5**). Siphon-seq reduced programs associated with translation and RNA-processing, consistent with removal of leukocyte RNA, and increased platelet aggregation, granule, and cytoskeletal programs, whereas immune and inflammatory sets were significant only in contaminated data. Contamination therefore produces reproducible distortions of inferred platelet programs, adding immune-related signal while obscuring platelet-specific programs.

Deviations in the platelet transcriptome were reflected at the protein level. Genes most depleted relative to megakaryocyte expression, half-life and localization were less often detected in two platelet proteome datasets (OR 0.65–0.72), and the most enriched were more often detected (OR 1.78–1.94; **Figure 2H, Supplemental Figure 6D–G, Supplemental Data 6**). Thus, transcripts enriched beyond expectation were also more often present as protein. As a functional confirmation of this interpretation, we mapped approved antithrombotic drug targets onto the core (**Figure 2I**; **Supplemental Data 7**).^43–45^ Primary targets of most antiplatelet drug classes were core transcripts. In contrast, only 1 of 13 targets of anticoagulant and thrombolytic drugs was a core transcript.^46–48^ These drugs act on plasma proteins made mainly by the liver and endothelium rather than on platelets, so their targets would not be expected in platelets. The core transcriptome therefore captures targets of drugs that act directly on platelets.

### Siphon-seq extends to clinical samples in a disease context

Having established Siphon-seq in healthy donors using live cells, we next asked whether it could extend to a clinical setting where cell fixation is preferable. We therefore applied Siphon-seq in a pilot study using 10x Flex fixed RNA chemistry in platelets from four sepsis and four non-sepsis control donors (**Fig. 3A, Supplemental Table 3).** Two heavily contaminated samples (one sepsis, one control) yielded too few pure platelet GEMs for purified analysis, leaving six matched pairs. Matched-pair analysis confirmed platelet marker retention and contamination marker reduction, mirroring the 3′ dataset **(Fig. 3B-C)**. By MuSiC deconvolution, estimated platelet RNA in pre- purification libraries ranged from 6.7% to 100% (**Fig. 3D**) and after Siphon-seq ranged from 98.7– 100% (**Fig. 3E**), and contaminant-marker transcripts fell a median 4.0-fold (**Fig. 3F**).

**Figure 3.**
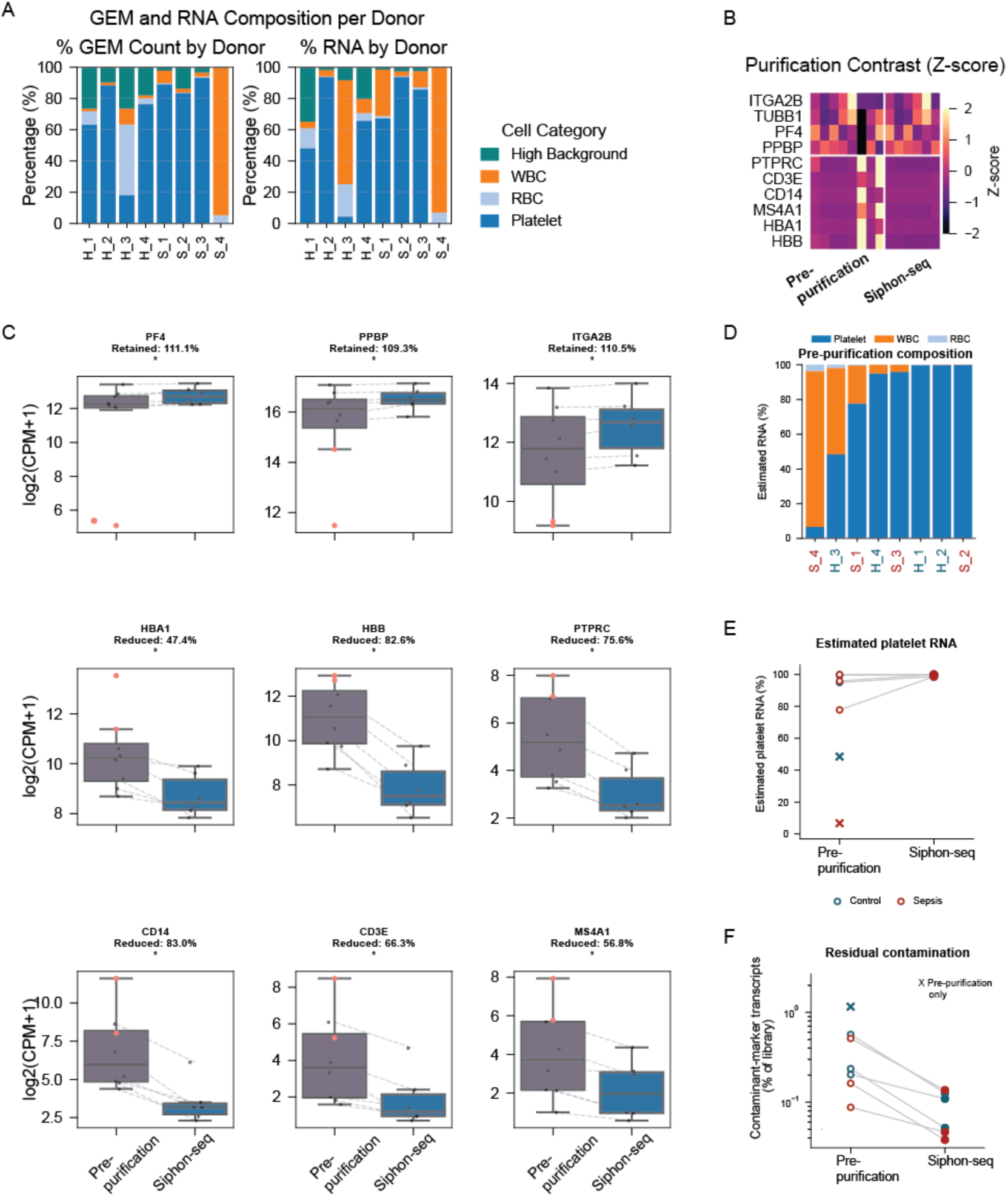
Siphon-seq purifies fixed clinical samples. **(A)** Per-donor composition of pre-purification preparations from 8 donors (4 sepsis, and 4 controls: 2 critically ill patients without sepsis and 2 healthy volunteers), by GEM count (left) and by RNA (right). Colors indicate platelet, erythrocyte (RBC), leukocyte (WBC) and high-background fractions. **(B)** Z-score heatmap of platelet and contaminant marker genes in pre-purification (left) and Siphon-seq (right) preparations. **(C)** Platelet markers (PF4, PPBP, ITGA2B; top row) and contaminant markers (HBA1, HBB, PTPRC, CD14, CD3E, MS4A1; middle and bottom rows) in pre-purification versus Siphon-seq preparations. Dashed lines connect the 6 donors with both preparations; unconnected points are donors with pre-purification data only. Red points, sepsis donors. Expression is log₂(CPM+1). Exact Wilcoxon signed-rank test on the 6 pairs, * P < 0.05. Percentages show mean retention (platelet markers) or reduction (contaminant markers) after purification. **(D)** Estimated RNA composition (MuSiC; Methods) of each pre-purification preparation, ordered by platelet RNA. Donor labels are colored by group (red, sepsis; teal, control). **(E)** Estimated platelet RNA (%) in pre-purification versus Siphon-seq preparations; ×, donors with pre-purification data only. **(F)** Residual contamination: the six contaminant markers in (C) as a percentage of the library (log scale); ×, as in (E).

### Leukocyte RNA alters pathway-level inference in sepsis

We performed GSEA separately on the pre-purification and Siphon-seq sepsis-versus-control comparisons and compared normalized enrichment scores (NES) for 1,624 GO biological processes (**Figure 4A, Supplemental Data 8**). By NES pre-purification and Siphon-seq, 743 processes were unattenuated, 320 processes were positively enriched in both preparations and amplified by purification, while 211 processes were negatively enriched in both preparations and amplified by purification. In contrast, 265 processes reversed from positive to negative and 85 from negative to positive. Given the pilot sample size, findings are hypothesis-generating, so we applied a permissive FDR threshold of 0.25: 157 processes passed before purification and 42 in Siphon-seq.

**Figure 4.**
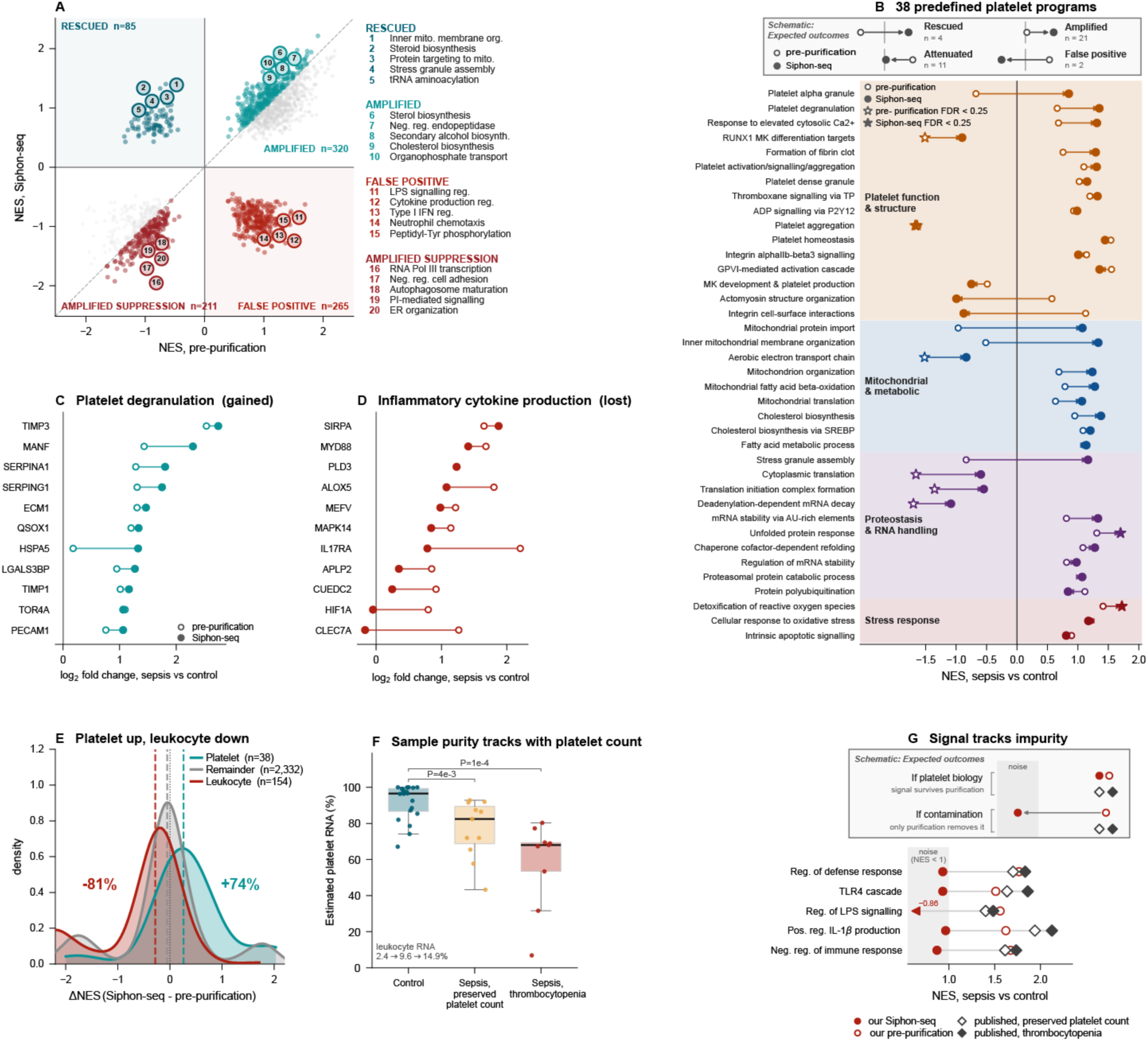
Siphon-seq changes which pathways appear regulated in sepsis. **(A)** Normalized enrichment scores (NES) for 1,624 GO biological processes, sepsis versus control, in pre-purification (x axis) and Siphon-seq (y axis). Shaded quadrants: rescued (enriched only after purification, n = 85) and false positive (enriched only before purification, n = 265). Twenty representative pathways, five per quadrant, are numbered and listed at right. **(B)** Thirty-eight platelet programs, grouped by theme. Open circles, pre-purification; filled circles, Siphon-seq; open and filled stars, FDR < 0.25 in the respective library. Twenty-eight of 38 programs moved upward in NES after purification (sign test P = 0.0051; Wilcoxon P = 8.2 × 10⁻⁴; median ΔNES +0.26). Schematic: rescued (n = 4), amplified (n = 21), attenuated (n = 11), false positive (n = 2). **(C, D)** Leading-edge genes of platelet degranulation (gains signal) and of regulation of cytokine production in inflammatory response (loses signal; pre-purification NES +2.10, FDR 0.003): log₂ fold change, sepsis versus control, in each library. **(E)** Change in NES after purification (Siphon-seq − pre-purification) for the platelet programs in (B), leukocyte-annotated pathways, and the 2,332 other tested pathways. Dashed lines, group medians. Both flanking groups differ from the remainder (Mann–Whitney P = 2 × 10⁻⁵ and 2 × 10⁻¹²). **(F)** Estimated platelet RNA (%) in a published sepsis cohort (GSE252275; 20 controls, 11 sepsis with preserved and 9 with low platelet count). P values versus control, Mann–Whitney test. **(G)** NES, sepsis versus control, for five immune pathways reported in that cohort: our pre-purification and Siphon-seq libraries, and the published cohort split by platelet count (median estimated platelet RNA: preserved, 82%; thrombocytopenic, 68%). Our pre-purification NES match the published cohort (1.51–1.77 vs 1.40–2.13); after purification, every pathway falls to noise (NES −0.86 to 0.96). Shading, NES < 1.

Pathways that reversed from positive to negative were dominated by inflammatory and leukocyte-associated processes, including regulation of lipopolysaccharide-mediated signaling, cytokine production, and neutrophil chemotaxis (NES +1.05 to +1.56 pre-purification; −0.86 to −1.23 after). Leukocyte RNA can therefore create an apparent inflammatory signal that is absent from purified platelet preparations. Regulation of lipopolysaccharide-mediated signaling was significant only before purification (FDR = 0.20), with the leukocyte genes LTF and LY96 among its leading-edge genes.

Among pathways amplified by purification, sterol and cholesterol terms formed the largest coherent family and the only ones crossing the FDR threshold. Sterol biosynthesis (NES +1.32 to +1.79) and secondary alcohol biosynthesis (+1.27 to +1.74) reached FDR < 0.25 only in Siphon-seq. The 85 pathways that became positive only in Siphon-seq included translation and RNA handling, protein homeostasis and mitochondrial programs, although none reached FDR < 0.25. For example, inner mitochondrial membrane organization shifted from NES −0.51 to +1.33, and 8 of its 15 leading-edge genes encode mitochondrial protein-import components. These results show that leukocyte RNA can both introduce inflammatory pathway signals and obscure programs that emerge only after purification.

We next tested whether 38 platelet processes gained enrichment (**Figure 4B, Supplemental Data 8**). Overall, 28 of 38 moved upward in NES (sign test, P = .0051; Wilcoxon, P = 8.2 × 10⁻⁴; median change, +0.26). Platelet degranulation, for example, increased from NES +0.66 pre- purification to +1.35 in Siphon-seq. In contrast, processes sharing genes with leukocytes, such as integrin cell-surface interactions (+1.13 to −0.87) and actomyosin structure organization (+0.57 to −0.99), decreased, whereas platelet aggregation remained negatively enriched (NES −1.64 to −1.67). The same pattern held at the gene level. Leading-edge genes of platelet degranulation, including the α-granule genes TIMP3 and SERPING1, had larger sepsis-versus-healthy log₂ fold changes after purification (**Figure 4C**). In contrast, leading-edge genes of cytokine production in inflammatory response, the most strongly enriched leukocyte pathway before purification (NES +2.10; FDR = .003), moved toward no difference (MYD88, ALOX5, IL17RA, MAPK14) or reversed direction (CLEC7A, HIF1A; **Figure 4D**).

Changes were specific to platelet and leukocyte pathways (**Figure 4E, Supplemental Data 8**). Platelet pathways became more enriched after purification (74% increased; median change, +0.26), leukocyte-associated pathways became less enriched (81% decreased; median, −0.28), and the remaining ∼2,332 pathways changed little (median, −0.05). Purification therefore strengthens platelet-associated and weakens leukocyte-associated signal rather than shifting pathway enrichment globally.

### An independent sepsis cohort shows the same contamination-sensitive immune programs

We next examined an independent, publicly available bulk platelet RNA-seq sepsis cohort containing 20 controls, 11 sepsis patients with preserved platelet counts, and 9 with low platelet counts (GSE252275, **Figure 4F, Supplemental Figure 10**, **Supplemental Data 9**). MuSiC deconvolution estimated that platelet RNA declined from a median of 96.5% in controls to 82.4% in sepsis with preserved platelet counts and 68.1% with low platelet counts (P = .004 and P = 1.5 × 10⁻⁴ vs controls, respectively), whereas leukocyte RNA rose from 2.4% to 9.6% and 14.9%.

Of 56 pathways enriched in our pre-purification data that returned toward the null after purification (a different set from the 38 platelet programs in Figure 4B and 4E), 23 were also enriched (FDR < 0.25) when we analyzed the deposited counts of that cohort without removing contaminating RNA, both overall and in each platelet-count stratum (**Supplemental Data 9**).

**Figure 4G** shows five non-redundant immune examples: regulation of defense response, the TLR4 cascade, regulation of lipopolysaccharide-mediated signaling, positive regulation of IL-1β production, and negative regulation of immune response. Their enrichment in our pre- purification libraries was comparable to that in the conventional analysis of the deposited counts (NES 1.51–1.77 vs 1.40–2.13), and for all 5, NES was higher in the low-than in the preserved-platelet-count stratum. After purification in our donors, all 5 returned toward the null (NES −0.86 to +0.96). Immune-pathway enrichment therefore tracked estimated leukocyte burden in the independent cohort and was lost when leukocyte RNA was removed in our paired samples.

## Discussion

Platelet transcriptomics has been limited by a tradeoff between sensitivity and purity. Bulk RNA-seq is sensitive but pools platelet and contaminating RNA into a single library, so it depends on highly pure preparations that may be inconsistent between control and experimental cohorts. Single-cell approaches resolve individual cells but recover too little platelet RNA for robust library generation across a variety of platforms. Siphon-seq bridges these approaches. Overloading single-cell partitions raises aggregate RNA input above the library-generation threshold, and partition-level barcoding allows nonplatelet partitions to be removed computationally after sequencing. The result combines the transcript recovery of a bulk preparation with partition-level identification of contaminating material. Because it does not require single-platelet RNA yields, Siphon-seq avoids the heterogeneous and often inconclusive results of previous single-cell platelet approaches, and its application to 34 healthy donors allowed us to define a consensus core platelet transcriptome.^28,43,49–51^

The consequences of contamination extend beyond obviously foreign transcripts. The largest contamination-driven differences were evenly split: among the 20 genes most enriched before purification, 10 were erythrocyte or leukocyte markers, which lineage-marker quality control would flag, and 10 were ribosomal protein transcripts, which it would not. Ribosomal transcripts overlap genuine platelet biology and are easily read as translational activity. Contamination also shaped inter-donor correlation structure, clustering donors by contamination profile rather than by platelet biology (**Figure 1K**). Bulk platelet RNA-seq should therefore be interpreted alongside an estimate of RNA-level purity.

With this confound removed, megakaryocyte expression was a major but incomplete determinant of the core platelet transcriptome. Relative to the megakaryocyte expectation, platelets were depleted of biosynthetic programs typical of a growing cell, including translation, tRNA aminoacylation, the mitochondrial matrix, and cholesterol biosynthesis. They were enriched for platelet activation, vesicle sorting, and protein catabolism (**Figure 2E**). Transcript half-life and nuclear-to-cytoplasmic localization did not explain these deviations, and the deviations were reflected at the protein level. Genes most enriched relative to expectation were more likely to be detected in platelet proteomes, and the most depleted genes were less likely (**Figure 2**). The core also contained the primary targets of most antiplatelet drug classes but few anticoagulant or thrombolytic targets, supporting its use as a reference for platelet biology. Most of this core was contained in earlier commonly expressed platelet gene sets (82% of core genes in Londin et al., 75% in Simon et al.), whereas genes unique to those sets were far more often contaminant-enriched in our data (26–29% versus 4%), suggesting that contamination has contributed to the composition of existing platelet reference lists (**Supplemental Data 10**).^52,53^ Together, these findings suggest that platelet RNA content reflects selective packaging, partitioning, or retention beyond passive inheritance of the megakaryocyte transcriptome. Megakaryocytes from bone marrow, lung, and spleen differ transcriptionally, so tissue of origin, as well as disease state, could contribute to these departures and to platelet heterogeneity.^54–56^

Sample purity determined how clearly these deviations were recovered. Moderate leukocyte contamination compressed gene-set deviations toward the megakaryocyte expectation without reducing the number of significant results (**Figure 2F**), and the gene sets that changed with purification in our donors changed in the same direction in an independent public cohort stratified by purity. Contamination can therefore create aberrant significance between cohorts that misrepresents platelet biology.

Contamination also altered biological conclusions in disease. In our pilot sepsis cohort, inflammatory and innate-immune pathways enriched before purification weakened or reversed after Siphon-seq, whereas platelet-associated programs generally gained signal. This was not a global loss of signal: leukocyte-associated pathways preferentially decreased, platelet-associated pathways preferentially increased, and most other pathways changed little. Purification also revealed or strengthened programs involving sterol biosynthesis, mitochondrial maintenance, RNA handling, and protein homeostasis. In an independent cohort (GSE252275), estimated platelet RNA was lower in sepsis and lowest among patients with low platelet counts. Of the 56 pathways that lost signal after purification (likely reflecting contamination signals), 23 were also enriched in sepsis in GSE252275, most strongly in the lower-purity stratum (**Figure 4G**). The observation that contamination tracks immune enrichment across cohorts, combined with the same signals being lost on purification within donors, supports the importance of leukocyte RNA as a contributor to pathway-level associations in bulk RNA platelet studies of disease.

The limitations of this study include the following: Siphon-seq reduces but does not eliminate contamination. Purified samples retained very low levels of leukocyte-marker RNA, and contaminating RNA within platelet-containing partitions, such as ambient RNA, cannot be removed by partition filtering. Siphon-seq profiles should therefore be regarded as substantially enriched for platelet RNA rather than absolutely pure. Second, because we used 3′ capture and probe-based chemistry, Siphon-seq detects only polyadenylated or probe-targeted transcripts and does not capture microRNAs or most other non-coding RNAs. Third, the small pilot sepsis cohort limits inference for individual pathways. Finally, MuSiC may have limited resolution at the highest platelet fractions.

In summary, Siphon-seq generates sensitive, computationally purified platelet transcriptomes. Application of Siphon-seq revealed that platelet transcript content departs systematically from megakaryocyte expression, that even modest leukocyte contamination obscures these departures, and that in a clinical comparison, contamination altered biological conclusions. RNA-level purity in platelet transcriptomics directly shapes biological inference.

## Acknowledgements

Sequencing was performed at the UCSF CAT, supported by UCSF PBBR, RRP IMIA, and NIH 1S10OD028511-01 grants. We acknowledge the *UCSF Parnassus Flow CoLab* (RRID:*SCR_018206*) for Flow cytometry instrument use and research reported here was supported in part by the DRC Center Grant NIH P30 DK063720. We acknowledge the HDFCCC Laboratory for Cell Analysis Shared Resource Facility for flow cytometry instrument use that was funded in part through a grant from NIH (P30CA082103). B.A.M. is a MacMillan Family Foundation Awardee of the Life Sciences Research Foundation. C.C. is supported by an IARS Mentored Research Award and the Deutsche Forschungsgemeinschaft (DFG) Research Fellowships (CO 2096/1-1). Q.Z. is supported by a Cancer Research Institute Immuno-Informatics Postdoctoral Fellowship (CRI5054). S.F.M. was supported by NIH K12 K12HD105250, NIH K08 K08HL177269, and the Parker B. Francis Foundation fellowship. C.S.C. was supported by R35HL177135. The study was funded by NIAID R01 AI165919 (M.R.L.) and NHLBI R35 HL161241 (M.R.L.).

## Author contributions

Conceptualization & Methodology: B.A.M., C.C., Q.Z., Z.J.G., M.R.L.; Software & Formal Analysis: B.A.M., C.C., Q.Z.; Investigation & Validation: S.F.M., L.Q.; Resources & Data Curation: B.A.M., C.C., C.S.C.; Writing (Draft & Review/Editing): B.A.M., C.C., Q.Z., S.F.M., L.Q., C.T., C.S.C., Z.J.G., M.R.L.; Visualization: B.A.M., C.C.; Supervision & Project Administration: Z.J.G., M.R.L.; Funding Acquisition: M.R.L.

