## Supplementary Information for "Revealing the platelet transcriptome using Siphon-seq: a sensitive, purity-aware method for RNA-sequencing"

Supplementary Methods

Supplementary Tables 1-5

Supplementary Figures 1-10

Supplementary Data 1-11

### Supplemental Methods

#### Platelet isolation

**Platelet-rich plasma.** Whole blood was transferred to 15-mL tubes and centrifuged at 300 × g for 10 minutes with acceleration and deceleration set to zero to isolate platelet-rich plasma (PRP). PRP was transferred to a fresh tube and supplemented with prostacyclin ($PGI_2$, 0.1 µg/mL) and apyrase (0.25 U/mL) to minimize platelet activation and aggregation. Platelets were pelleted at 1000 × g for 10 minutes, again with acceleration and deceleration set to zero.

**Washing and filtration.** The platelet pellet was resuspended gently in twice the initial PRP volume of Buffer A (10 mM HEPES, 140 mM NaCl, 5 mM KCl, 5 mM $KH_2PO_4$, 1 mM $MgSO_4$, 5.5 mM glucose, 0.35% BSA, pH 6.7–6.8) supplemented with $PGI_2$ (0.1 µg/mL) and apyrase (0.25 U/mL). The suspension was passed through a 10-µm filter (pluriStrainer, pluriSelect) to remove aggregates.

**Counting.** Platelets were quantified by flow cytometry using 4-µm CountBright Plus counting beads. Samples were stained with Alexa Fluor 488–conjugated anti-human CD41 antibody (clone HIP8; BioLegend), with unstained and buffer-only controls included. Acquisition used a BD LSR II, BD FACSCanto II, Attune NxT (Thermo Fisher) or BD LSRFortessa X-20 Dual Right SORP, with the forward-scatter (FSC) threshold set to 1,000 and FSC and side-scatter recorded with logarithmic amplification. Platelets were identified by their low FSC/SSC profile and CD41 positivity, and doublets and aggregates were excluded on FSC-A versus FSC-H pulse geometry. Samples were acquired at a low flow rate, with at least 50,000 platelet events recorded per sample.

**Fixation (sepsis cohort and matched controls).** Platelets were fixed in 20 mL of 4% paraformaldehyde in dPBS containing $PGI_2$ and apyrase at the concentrations above, for 1 hour at room temperature followed by overnight incubation at 4 °C. Cells were pelleted at 1,000 × g for 10 minutes and resuspended in 10x Quenching Buffer containing 10% Enhancer. Glycerol was added to a final concentration of 10%, and samples were stored at −80 °C.

#### Siphon-seq library preparation

**Overloading.** Standard 10x Chromium loading targets a cell number low enough to make multiple captures per partition unlikely. Siphon-seq inverts this: platelets were loaded far above that number so that each GEM captured an ensemble of many platelets, raising per-partition RNA input above the threshold below which adapter-dimer products dominate over cDNA, while preserving an independent barcode per partition. Platelets were loaded at $5 \times 10^6$ per channel for Chromium Next GEM Single Cell 3′ v3.1 and $1 \times 10^7$ per channel for GEM-X Single Cell 3′ v4.

**Live-cell libraries (3′ chemistry).** The required number of platelets was transferred to a 1.5-mL microcentrifuge tube containing 500 µL Buffer A and centrifuged at 1,000 × g for 10 minutes. The supernatant was removed carefully, leaving a residual volume below 43.2 µL for Next GEM Single Cell 3′ v3.1 or below 36.6 µL for GEM-X Single Cell 3′ v4. Samples were processed immediately for GEM generation and library preparation according to the corresponding 10x Genomics protocol. Of the 34 healthy donors, 22 were run on Next GEM 3′ v3.1 and 12 on GEM-X 3′ v4.

**Fixed libraries (probe-based chemistry).** Fixed samples were thawed at room temperature, pelleted at 1,200 × g for 10 minutes and resuspended in Quenching Buffer. Platelet concentration was determined by flow cytometry with counting beads as above. Probe hybridization and library preparation followed the 10x protocol with the following exceptions: $20 \times 10^6$ cells were used for the initial probe hybridization, all centrifugation steps were performed at 1,200 × g for 10 minutes, an additional probe wash was performed to ensure clearance of unbound probe, and $5 \times 10^6$ platelets were loaded per 10x lane.

**Sequencing.** Libraries were sequenced to a median of 199 million reads (range 13–540 million), with a median sequencing saturation of 87.9% (range 57.5–98.4%). Per-library metrics are given in Data file 2.

#### Read processing

Libraries were processed with Cell Ranger 8.0.1 (cellranger count, default settings) against the 10x GRCh38-2024-A reference, which counts intronic reads by default.

#### Partition classification

**GEM inclusion.** Cell Ranger's cell calling assumes one cell per GEM and does not apply to overloaded platelet GEMs, so GEMs were taken from each library's raw feature-barcode matrix. For each library, the distribution of UMIs per barcode was binned in 5-UMI bins, and the inclusion threshold was set at the first bin in which the histogram's slope turned positive, i.e. the trough separating empty droplets from GEMs. Thresholds ranged from 7.5 to 42.5 UMIs (median 12.5). This gave 3,101,161 GEMs across the 34 donor libraries.

**Clustering and training labels.** Using scanpy, raw counts were normalized to the median library size and log-transformed.[1] Then 2,000 highly variable genes were selected with the donor library as the batch (batch_key = donor), followed by principal component analysis (50 components, unscaled data), a 15-nearest-neighbor graph, UMAP and Leiden clustering (resolution 1; 18 clusters).[2,3] Clusters were labeled by their Wilcoxon marker genes as erythrocyte (one cluster: HBB, HBA1, HBA2), leukocyte (four clusters: LYZ, immunoglobulin, PTPRC or S100A9 markers) or platelet (six clusters: PF4, PPBP, NRGN, GP1BB, or predominantly mitochondrial transcripts).

**Classifier.** A CONCORD classifier (concord_sc 0.9.3) was trained on a balanced set of 50,000 GEMs per class drawn from the labeled clusters (erythrocyte GEMs, of which 23,824 were available, were drawn with replacement).[4] Input features were the union of the top 500 Wilcoxon marker genes for each class against the other two on the balanced set (1,387 genes). The model used the donor library as the domain, a 32-dimensional latent space, a 128-unit encoder, 90% of the balanced set for training and 10% for validation, 10 epochs, a batch size of 64 and a learning rate of 0.01, with all other settings at their defaults. The trained model was applied to all GEMs, giving each a platelet, erythrocyte or leukocyte call and the posterior probability of each class.

**High background.** For each GEM, the contaminant posterior was the larger of its erythrocyte and leukocyte posteriors. A platelet-called GEM was classed as high background, and excluded, if its contaminant posterior exceeded 0.2 or it belonged to a Leiden cluster in which more than 50% of all GEMs exceeded 0.2. Six clusters (7, 11, 14, 15, 16, 17) met the second criterion when the rate was computed over all GEMs, as in Supplemental Figure 2C; four of them (7, 11, 15, 17) also met it when computed over platelet-called GEMs only, the rule that gave the retained GEM count below, and the two rules differ in 371 of the 2.21 million retained GEMs. Across the 34 donors this left 2,214,926 platelet GEMs.

**Choice of cutoff.** Contamination-marker RNA in platelet-called GEMs was summarized in bins of contaminant posterior, and in the GEMs retained at cutoffs from 0.1 to 0.5. At 0.2, retained GEMs carried about 23-fold less globin RNA and about 14-fold less leukocyte-marker RNA (marker panels of Supplemental Figure 2G, pooled across donors) than the unpurified library. The 4.7-fold and 24-fold differences given below are computed differently, as per-donor median CPM of the full libraries. Leukocyte RNA was removed mainly by the classifier's leukocyte calls rather than by the cutoff.

**Pseudobulk profiles.** For each donor, raw UMI counts were summed over all GEMs ("pre-purification", equivalent to a conventional platelet preparation) or over retained platelet GEMs ("Siphon-seq").

#### Pseudobulk construction and normalization

For each donor, two pseudobulk libraries were built by summing raw UMI counts across GEMs: pre-purification, over every GEM that passed the UMI floor, which is the library a conventional platelet preparation sequenced on this platform would give; and Siphon-seq, over the platelet-called GEMs left after high-background GEMs were removed. The same two definitions were used for the fixed-RNA (Flex) pilot cohort, and the identical terms are used in every figure.

Two normalizations are used, and each analysis states which. Counts per million scaled by trimmed mean of M-values (TMM, edgeR calcNormFactors) is used wherever genes are compared across donors on one preparation.[5,6] Counts per million by library size alone, with no TMM factor, is used for the paired pre-purification versus Siphon-seq comparisons, because purification removes a large and variable share of each library's RNA, which violates the assumption behind TMM that most genes are unchanged.

#### Deconvolution of bulk libraries

**Reference.** Cell-type RNA fractions were estimated with MuSiC (v1.0.0) against a six-type pseudobulk reference.[7] Five types: platelet, neutrophil, T/NK, B/plasma and myeloid were taken from the 14 healthy-donor libraries of GSE163668 (GSM4995449–GSM4995462; Combes et al.)[8,9]. Raw 10x matrices were reprocessed per library with SCTransform normalization, doublet removal (DoubletFinder) and the original study's quality thresholds, then merged, batch-corrected across donors with Harmony and clustered (Louvain, resolution 0.6); clusters were labeled by module-score argmax against the original study's marker panels and reviewed against raw marker expression.[10,11] Against the original study's own per-cell labels (21,292 cells matched by barcode), agreement was 91.5–98.9% for each of the five types used. Erythrocytes were taken from a separate dataset, because whole blood in GSE163668 was red-cell-lysed before capture and its erythrocyte-labeled cluster proved to be mostly neutrophils carrying ambient hemoglobin RNA: the day-1 sample of GSE184916 (GSM5600335; Jain et al.), using the authors' own filtered barcodes without a nucleated-cell complexity filter, in which hemoglobin transcripts account for 97.5% of UMIs.[9] MuSiC requires at least two samples per cell type to estimate cross-subject variance, so these 5,000 cells were split at random into two halves and pseudobulked separately; the two profiles are pseudo-replicates of one sample, not biological replicates. Pseudobulk profiles were built by summing raw counts per donor and cell type (all 70 donor × type combinations present, none below 20 cells), giving a reference of 72 profiles over the 10,132 genes shared with the erythrocyte data.

**Fitting.** MuSiC was run as music_prop(bulk, reference, clusters = "cellType", samples = "sampleID") on raw counts with all other arguments at their defaults, so no marker list was supplied and all six types were fitted. No gene-length correction was applied to the reference. Samples with fewer than 1,000 total counts were excluded as degenerate.

**Conversion to RNA share.** With a pseudobulk reference, MuSiC's fitted proportion is not a fraction of RNA, because one reference unit of each type carries a different amount of RNA (relative to platelet: neutrophil 0.66, B/plasma 5.5, erythrocyte 12.8, myeloid 14.1, T/NK 26.3). Each type's share of RNA was therefore taken as its share of the fitted expression, $p_k \times \Sigma(D_k) \div \Sigma_j p_j \times \Sigma(D_j)$, where p is the fitted proportion and $\Sigma(D_k)$ is the column sum of the reference profile for type k. This reproduces MuSiC's own fitted contributions to within $4 \times 10^{-16}$. This study reports this quantity as "Estimated platelet RNA (%)".

**Validation.** The reference was rebuilt in four donor folds (4, 4, 3 and 3 donors held out), each fold using only its training donors; the erythrocyte component was split so that the reference and the mixtures came from different halves of the source sample. Held-out platelet profiles were mixed in count space with held-out contaminant profiles at 13 true platelet RNA shares from 0.30 to 1.00, with 14 donors × 5 multinomial draws per point, over the 9,748 genes fitted in production. The contaminant was each of the five types alone or a PlateletBase-like mixture (neutrophil 0.43, T/NK 0.21, erythrocyte 0.20, myeloid 0.12, B/plasma 0.03).[12] Mixtures were drawn at 3,799,040 counts (the median PlateletBase library) and at 813,839 counts (its 10th percentile). Each mixture was fitted with the production call and converted to RNA share with the same code. Between 85% and 100% true platelet RNA the root-mean-square error was 0.05–0.06 with a bias of −0.01 to −0.04; pure platelet mixtures read 96.1% on average (95% range 87.5–100%). Error was set by the held-out donor rather than by depth: the within-donor spread was below 0.003 and the error changed by at most 0.003 between the two depths. The smallest contaminant share at which at least 95% of mixtures were estimated to hold more than 1% non-platelet RNA was 2% for erythrocytes, 7.5% for neutrophils and the mixture, 10% for myeloid and B/plasma cells and 15% for T/NK cells. Values above about 90% platelet RNA are therefore not distinguishable from pure platelet RNA, while values below about 85% indicate contamination. Because the mixtures are built from 10x pseudobulks, the simulation does not test the difference between the 10x reference and full-length bulk libraries, and erythrocyte accuracy is likely optimistic because the red cells come from a single sample.

**Public bulk libraries.** Gene-level raw counts for 3,711 samples in 15 projects were obtained from PlateletBase (plateletbase.clinlabomics.org.cn).[12] Each project was processed independently, keeping its own gene set; the ComBat-corrected log2 matrix distributed with the resource was not used, because batch there is confounded with isolation protocol and therefore with contamination itself.[13] Unversioned Ensembl IDs were mapped to the reference's symbols through the GSE163668 features.tsv.gz file, with counts summed where several IDs mapped to one symbol and genes absent from a project excluded for that project only. Project 04 (n = 20, median depth about 54,000 reads) was excluded due to lowe sequencing depth, leaving 3,689 samples in 14 projects. Leukocyte-depleted or FACS-sorted isolation was determined by reading each project's original publication. Project 15 has hemoglobin genes removed upstream, so its erythrocyte share is zero by construction. Several of the largest cohorts share a thromboSeq lineage whose quality control excludes contaminated libraries, so their contamination distributions are truncated at an undocumented threshold (Supplemental Table 5).

**Our own libraries and the pilot.** Pre-purification and Siphon-seq pseudobulk libraries were fitted with the same call and converted the same way. For the external sepsis cohort GSE252275, published counts were used as deposited.

#### The core platelet transcriptome

All 38,606 quantified genes were scored on the 34 Siphon-seq libraries using TMM-CPM. A gene is in the **core** if its mean expression is at least 2 CPM and it exceeds 2 CPM in at least 90% of donors (31 of 34); **contextual** if its mean reaches 2 CPM but it is above that level in fewer than 90% of donors, that is, real signal in some donors but not a consistent feature of the transcriptome; and **below threshold** otherwise. This gives 3,436 core, 2,644 contextual and 32,526 below-threshold genes. At this threshold the two criteria are nested, so any gene above 2 CPM in 90% of donors is in the core by definition. Scoring presence instead at a single read identifies 3,163 genes present in at least 90% of donors that still fail the core filter, and these were used where a consistency-matched comparison group was needed.

Two sensitivity analyses bound the definition. First, prevalence was re-scored at a single raw read and at 0.25, 0.5 and 1 CPM with the 90% rule unchanged; all 3,436 core genes pass at every cut. Second, the whole pipeline was repeated on the matched pre-purification libraries of the same 34 donors. The pre-purification libraries, which carry 4.7 times more leukocyte-marker RNA and 24 times more hemoglobin RNA, give a core of 5,487 genes that overlaps ours at a Jaccard index of 0.58, so the definition is reproducible only at the purity Siphon-seq provides.

**Cecal slurry sepsis and blood collection**. Cecal slurry was prepared as previously described.[14,15] Briefly, cecal contents from C57BL/6J mice were suspended in PBS and 10% glycerol (1 mL per 100 mg cecal content) and sequentially filtered through sterile meshes. Sepsis was induced by intraperitoneal injection of 200 µL cecal slurry; control mice received an equal volume of vehicle. Peripheral blood was collected at the indicated time points by retro-orbital bleeding into MiniCollect $K_2EDTA$ tubes (Ref. 450532). Tubes were gently inverted to prevent clotting, and blood was processed immediately for downstream analysis. Blood from the three mice in each arm was pooled, giving one Siphon-seq library per arm.

**The megakaryocyte expectation model**

**Covariates.** Megakaryocyte expression is from BLUEPRINT: RSEM gene quantifications for six healthy cord-blood megakaryocyte donors (with 13 neutrophil and 8 classical monocyte donors used elsewhere), summarized per gene as log2(median FPKM + 1) and mapped from Ensembl IDs to current symbols.[16,17] Classes of RNA that 10x 3′ chemistry cannot see (small nuclear, small nucleolar, vault, 7S and microRNA genes, and tRNAs) were removed. mRNA half-life is the consensus human estimate of Agarwal and Kelley (2022).[18] The nuclear-to-cytoplasmic ratio was computed from ENCODE polyA RNA-seq of matched nuclear and cytosolic fractions of K562 and HepG2 (two replicates each) as log2((nuclear TPM + 0.1)/(cytosolic TPM + 0.1)), averaged over the two cell lines.[19] Genes with megakaryocyte expression above 1 log2 FPKM and values for both covariates form the model universe of 9,616 genes, of which 2,789 are core and 1,627 contextual.

**Core membership.** Logistic regression of core membership on megakaryocyte expression gives a McFadden pseudo-$R^2$ of 0.208, with an odds ratio of 2.02 per log2 FPKM unit. Adding half-life changes the pseudo-$R^2$ by less than 0.0001 (likelihood-ratio $P = .53$), so half-life carries no information about which transcripts persist; adding the nuclear-to-cytoplasmic ratio raises it to 0.213 ($P = 3.6 \times 10^{-13}$) and adding five functional indicators to 0.224. The megakaryocyte-only model is used as the expectation, and it agrees with the fuller model (per-set Pearson $r = 0.998$, same direction for every set called by both).

**Gene-set test.** Gene sets were taken from GO Biological Process 2023, GO Cellular Component 2023 and Reactome 2022, restricted to sets with at least 20 genes in the model universe (2,206 sets).[20-22] For each set the expected number of core genes is the sum of its members' fitted probabilities under the passive model, and the observed count is compared with it as $z = (\text{observed} - \text{expected})/\sqrt{\Sigma p(1-p)}$, the Poisson-binomial standard deviation, with a two-sided normal P value and Benjamini-Hochberg control across sets.[23] Against megakaryocyte expression alone, 455 sets differ from expectation (269 enriched, 186 depleted); against the full model, 500 (312 enriched, 188 depleted). The spread of observed-over-expected values is 3.8 times that of simulations in which each gene enters the core independently with its fitted probability, so the departures exceed sampling noise. The same machinery applied to contextual versus below-threshold genes gives 149 of 1,458 sets.

**Transcript amount.** For the 9,570 genes with any platelet reads, log2(mean CPM + 0.1) was regressed on megakaryocyte expression, half-life and the nuclear-to-cytoplasmic ratio ($R^2 = 0.366$). The residual of this fit is the platelet-enrichment axis: one unit is a twofold departure from the transcript level the three covariates predict. Sets were scored by their mean residual, with z computed against the residual standard error and Benjamini-Hochberg control, and a null built by permuting residuals across genes.

#### Purity comparisons

**Leukocyte-marker metric.** Contamination is quantified throughout as the summed library-size CPM of eight leukocyte genes (PTPRC, LYZ, S100A8, S100A9, CD52, CORO1A, LCP1, CD74), reported as the median across a dataset's samples. Companion metrics use hemoglobin genes (HBB, HBA1, HBA2, HBD) and platelet genes (PF4, PPBP, GP9, ITGA2B). On this scale our Siphon-seq libraries are at 153 CPM, our matched pre-purification libraries at 717, the high- and low-purity halves of the public cohort P13 (PRJNA353588) at 478 and 2,387, and those of P14 (PRJNA659491) at 472 and 3,420, a 22-fold range.

**Public comparison cohorts.** Two PlateletBase projects have enough healthy samples to split by purity, P13 (PRJNA353588) and P14 (PRJNA659491). In each, healthy samples were ranked by their MuSiC platelet proportion, one sample per donor, and split into a high-purity and a low-purity half of equal size: 110 samples each for P13 (median platelet proportion 1.000 and 0.680) and 160 each for P14 (1.000 and 0.507). The two projects share many of the same libraries, so they are not independent cohorts. Each dataset was called independently with the same rules, and, because the cohorts differ in size, each half was also scored as 20 random draws of 34 samples. Analyses were restricted to the genes quantified in every dataset, and the expectation model was refitted inside each dataset. Gene sets scoreable in all six datasets (our two preparations and the four public halves) number 2,194. Of these, our Siphon-seq data call 491 significant, our pre-purification libraries 510 and the high-purity halves 491 (P13) and 494 (P14), so these four datasets are comparably powered; the low-purity halves call 78 (P13) and 18 (P14) and are treated as underpowered throughout. The 149 purified-only sets are those significant in the Siphon-seq data and in none of the three moderately contaminated datasets (our pre-purification libraries and the two high-purity halves).

**What purification changes.** Each sample's score for a gene set is the mean of log2(CPM + 1) over its members minus that sample's mean over all universe genes, so the score is competitive and per-sample normalization cancels. For our own donors the paired difference between the two preparations was tested with a one-sample t-test over the 34 donors; for the public cohort the two halves were compared with Welch's t-test. Because broadly expressed genes dominate annotated sets, the median set does not sit at zero, and all changes are reported relative to the median set. A set was called changed at FDR < 0.05 with a centered change of at least 0.25 log2 (about 20%), with the cutoff varied between 0.15 and 0.5 as a sensitivity check. Overlap between the two datasets' calls was tested with Fisher's exact test against the 2,194 sets and drawn as area-proportional Venn diagrams. Set families were defined both by keyword and by database hierarchy (Reactome top-level pathway; GO slim category); shading is log2 of each family's count over its expected count from the family's share of all sets, starred at Benjamini-Hochberg q < 0.05.

**In silico titration.** To separate contamination from cohort and platform differences, each donor's Siphon-seq and pre-purification libraries were mixed in TMM-CPM space as $(1 - a) \times$ Siphon-seq + $a \times$ pre-purification, for a = 0, 0.1, 0.2, 0.3, 0.5, 0.7 and 1, so that donor, platform and processing are held fixed and only contamination varies. The core and all set-level readouts were recomputed at each step and compared with a = 0 by correlation, by the slope of the regression of mixed on unmixed effects (the attenuation of true signal), and by the share of the original calls retained; the x-axis is the mixture's own leukocyte-marker CPM. As a check, a = 0 reproduces the core gene list exactly and a = 1 the pre-purification calls. A noise floor for these curves was built by splitting the 34 Siphon-seq donors into two halves of 17 within sequencing batch (20 random splits), and the high-purity public half into two halves of 80, giving the change expected between two equally pure datasets.

#### Protein detection

Platelet protein detection was taken from two sources. Burkhart et al. (2012) supplemental data give copy numbers for 3,499 proteins; accessions were converted to gene symbols through an org.Hs.eg.db mapping with a published UniProt-to-GeneCards table as fallback, resolved to current symbols through an alias table, and protein groups collapsed to the highest copy number per symbol.[24,25] PaxDb v6.1's integrated platelet dataset, a weighted average of eight platelet proteomes, supplies the second, deeper list.[26] Detection is treated as binary presence in each resource. Among the 9,471 protein-coding genes of the model universe with an enrichment residual, 3,125 (33%) are detected by Burkhart et al. and 5,375 (57%) by PaxDb.

Expected detection was modeled by logistic regression of detection on a 4-degree-of-freedom B-spline in megakaryocyte expression plus half-life and the nuclear-to-cytoplasmic ratio, with no platelet measurement in the model (McFadden pseudo-$R^2$ 0.138 for Burkhart et al. and 0.109 for PaxDb). Genes were divided into deciles of the enrichment residual and summarized, per decile, as the ratio of the odds of observed detection to the odds of expected detection, so that both proteomes share one axis and 1 means detection exactly as predicted; the 95% confidence interval is the 2.5th to 97.5th percentile of 2,000 bootstrap resamples of the genes within each decile. The calibration of the expected probabilities was checked against observed detection. At the gene level, each twofold increase in enrichment raises the odds of protein detection by 1.19 (95% CI 1.17–1.21) for Burkhart et al. and 1.16 (1.13–1.18) for PaxDb. Two partial controls address the fact that enrichment and expected detection share the same megakaryocyte reference: the effect holds within strata of megakaryocyte expression, and it is unchanged when a 4-degree-of-freedom spline in log2 protein length (GENCODE v47 canonical) is added to the model (odds ratios 1.20 and 1.15), so it is not an artifact of mass-spectrometry detectability.[27]

#### Antithrombotic drug targets

Approved drugs (ChEMBL max_phase 4) in WHO ATC class B01A were retrieved from the ChEMBL REST API across its seven level-4 classes: vitamin K antagonists (B01AA), the heparin group (B01AB), antiplatelet agents (B01AC), fibrinolytic enzymes (B01AD), direct thrombin inhibitors (B01AE), direct factor Xa inhibitors (B01AF) and other antithrombotics (B01AX). For each drug, mechanism records gave its targets, and each target's gene symbols were taken from the ChEMBL target components' gene-symbol synonyms and resolved to current symbols; no activity or potency cutoff was applied, since only target identity was of interest. Nine drugs whose mechanism ChEMBL records against a salt or prodrug form rather than the parent molecule were added by hand from those same mechanism records (clopidogrel, prasugrel and ticlopidine to P2RY12; tirofiban to ITGA2B and ITGB3; vorapaxar to F2R; warfarin to VKORC1; dabigatran etexilate to F2; edoxaban to F10; fondaparinux to SERPINC1). This gives 48 drugs and 48 distinct targets in 80 drug-target pairs. Each target was then classified by the core rule on the Siphon-seq transcriptome, and Supplemental Data 7 reports, per drug class, how many of its targets are in the core. Full drug-target pairs, with DrugBank and ChEMBL identifiers, are in Data file 7.

#### Comparison with earlier platelet gene sets (Data file 10)

The core transcriptome was compared with two earlier catalogs of commonly expressed platelet genes: Londin et al. (2014), 5,592 protein-coding genes expressed above 1/10,000 of ACTB in all 10 donors, and Simon et al. (2014), 5,911 uniquely mapped mRNAs from 154 donors on microarray. Identifiers in both sets were resolved to current HGNC symbols by Ensembl ID first and then through an alias table, so that annotation drift did not create false mismatches. For every gene in either set, membership of our core, contextual and below-threshold classes was recorded, together with its paired pre-purification versus Siphon-seq fold change in our own 34 matched libraries (edgeR); a gene was called contaminant-enriched if it was higher before purification at FDR < 0.05 and |log2 fold change| > 1. Of the Londin set, 2,833 genes are in our core and 2,702 are not; of those outside it, 28.9% are contaminant-enriched in our data against 4.3% of the shared genes, and their median fold change is +0.46 log2. The Simon set behaves the same way (26.1% against 4.0%). Genes in our core but in neither earlier set (603 and 873) show no such excess (6.3% and 6.8%). Expression ranks were also compared directly: each earlier catalog's abundance ranking correlates with ours at ρ = 0.76, and for the Londin set the genes we call contaminant-enriched sit below the rank their position in that catalog predicts (median residual −8 percentile points, $P = 3.5 \times 10^{-18}$), while the same genes sit well above it when our own pre-purification libraries replace the Siphon-seq ones (+26 points; positive control). Full per-gene results are in Data file 10.

#### Differential expression and pathway analysis

**Pre-purification versus Siphon-seq genes.** The two libraries of each of the 34 donors were compared gene by gene with a paired Wilcoxon signed-rank test on log(CPM + 1), using library-size CPM, with Benjamini-Hochberg control across all tested genes. Testing was restricted to the 4,558 genes annotated as protein-coding in GENCODE v47; 3,385 were higher before purification and 678 higher after purification at FDR < 0.05. The 20 genes most increased in each direction were selected for display, ranked by mean paired log2 fold change, among genes that were also above 2 CPM in at least 90% of donors in the preparation where they were higher. Genes were labeled by category under a fixed rule: ribosomal-protein genes by gene family; otherwise the cell type with the highest median expression in the deconvolution reference, matching genes to the reference by Ensembl ID as well as by symbol because the reference carries pre-2020 symbols; and for the genes absent from the reference, platelet and megakaryocyte if megakaryocyte expression in BLUEPRINT was at least 1 log2 FPKM and above both neutrophils and monocytes, or if the protein is detected in platelets (Burkhart et al. or PaxDb), and otherwise "other". Values are plotted as the z-score of log(CPM + 1) per gene across both preparations of all donors, clipped at ±2.5.

**Sepsis pilot.** The pre-purification comparison uses all 8 pilot donors (4 sepsis, 4 control) and the Siphon-seq comparison the 6 donors with enough platelet GEMs after purification (3 and 3). Genes were kept if they reached 2 CPM on average across the pre-purification libraries, 2 CPM in at least two Siphon-seq libraries, and 10 counts in at least three donors of the preparation being tested. Sepsis was compared with control in each preparation using pyDESeq2 (design ~group, Wald test, α = 0.1, Cook's refitting, no log-fold-change shrinkage), and genes were ranked by the unshrunken Wald statistic.[28] Preranked gene set enrichment analysis was run with gseapy against GO Biological Process 2023 and Reactome 2022, with sets of 15–500 genes, 10,000 permutations and seed 42, and pathways were called at FDR < 0.25.[29] The two preparations were then compared pathway by pathway over the 1,624 GO Biological Process terms testable in both. A pathway was classed as **rescued** if its normalized enrichment score was at or below zero before purification and positive after it; as a **false positive** if the reverse; and otherwise, among pathways positive before purification, as **amplified** if the score rose and **suppressed** if it fell, with the mirror-image classification among pathways negative in both. This gave 85 rescued and 265 false-positive pathways. A predefined panel of 38 platelet programs, specified before the comparison was run from Reactome 2022 (22 programs), GO Biological Process 2023 (14) and GO Cellular Component 2023 (2: platelet alpha granule and platelet dense granule), was tested the same way with a minimum set size of 8, because canonical platelet sets such as GP1b-IX-V and P2Y12 signaling have fewer than 15 testable genes; 28 of the 38 shift toward positive enrichment after purification (sign test P = .005, median change +0.26).

**External sepsis cohort.** Gene-level counts for GSE252275 (20 control, 11 sepsis with preserved platelet count, 9 sepsis with thrombocytopenia) were downloaded from GEO, flattened from the per-sample worksheets of the deposited workbook into one matrix, and reconciled onto the reference's symbol space (10,124 of the reference's 10,132 genes recovered). Cell-type RNA shares were estimated with the same MuSiC call used everywhere else in this study, and myeloid, T/NK and B/plasma shares were summed for presentation as one leukocyte share. Groups were compared with two-sided Mann-Whitney tests and across all three groups with the Kruskal-Wallis test. Median platelet RNA was 96.5% in control, 82.4% with preserved platelet count and 68.1% with thrombocytopenia (P = .004 and $P = 1.5 \times 10^{-4}$ against control), and median leukocyte RNA 2.4%, 9.6% and 14.9%. Sepsis versus control differential expression was computed on the deposited counts with edgeR (filterByExpr, TMM normalization, robust dispersion estimation and a quasi-likelihood F test), genes were ranked by log fold change × −log10 P, and preranked gene set enrichment analysis was run against the same two libraries with sets of 15–500 genes and 1,000 permutations (FDR < 0.25). Leading-edge genes of each enriched pathway were classified with BLUEPRINT lineage calls, with myeloid-specific defined as expressed in neutrophils or monocytes and not in megakaryocytes. The two sepsis strata were also compared with control separately, and the difference in normalized enrichment score between them was compared between leukocyte-identity pathways and all others with a Mann-Whitney test.

**Mice and murine sepsis model**

Adult C57BL/6 (B6) mice were purchased from The Jackson Laboratory. All animal procedures were approved by the Institutional Animal Care and Use Committee of the University of California, San Francisco. Cecal slurry was prepared as previously described.[14,15] Briefly, cecal contents from C57BL/6J mice were suspended in phosphate-buffered saline (PBS) containing 10% glycerol at a ratio of 1 mL per 100 mg of cecal contents and sequentially filtered through sterile meshes. Sepsis was induced by intraperitoneal injection of 200 µL of cecal slurry; control mice received an equal volume of saline. Peripheral blood was collected at the indicated time points by retro-orbital bleeding into MiniCollect $K_2$EDTA tubes. Tubes were gently inverted to prevent clotting, and blood samples were processed immediately for downstream analyses.

#### Statistics, software, and data and code availability

**Statistics.** All tests are two-sided. Paired comparisons of a donor's pre-purification and Siphon-seq libraries use the Wilcoxon signed-rank test, or a paired t-test where stated; unpaired comparisons use the Mann-Whitney U test or Welch's t-test as stated in each legend. Correlations are Spearman's ρ unless Pearson's r is named. Multiple testing within a family of tests (genes, gene sets or pathways) is controlled with the Benjamini-Hochberg false discovery rate, and the family is stated in each legend. Box plots show the median and interquartile range, with whiskers to the range unless stated otherwise. The mouse experiment, with one pooled library per arm, is descriptive, and any P values or FDRs reported for it depend on an assumed dispersion and are not inferential.

### Supplemental Tables

#### Supplemental Table 1. Single-cell RNA-seq protocols tested on platelets before Siphon-seq

| Platform | Chemistry or kit | Capture | Platelet input and loading | Outcome | Notes |
|---|---|---|---|---|---|
| 10x Genomics | Chromium Next GEM Single Cell 3' v3.1 | poly(A) tail (transcript end) | Maximum loading per the manufacturer's recommendation, singleplex | No cDNA; reamplification negative | Later used for Siphon-seq, which succeeds only far above this maximum. |
| 10x Genomics | Chromium GEM-X Single Cell 3' v4 | poly(A) tail (transcript end) | Maximum loading per the manufacturer's recommendation, singleplex | No cDNA; reamplification negative | As above; the second Siphon-seq chemistry. |
| 10x Genomics | Chromium Next GEM Single Cell 5' v2 | 5' end | Maximum loading per the manufacturer's recommendation, singleplex | No cDNA; reamplification negative | |
| 10x Genomics | Flex Gene Expression (fixed RNA) | Probe-based | Maximum loading per the manufacturer's recommendation, singleplex | No cDNA; reamplification negative | Later used for the fixed sepsis cohort, again only above this maximum. |
| Parse Biosciences | Evercode Whole Transcriptome WT Mini | Split pooling | Maximum loading per the manufacturer's recommendation, singleplex | No cDNA; reamplification negative | |

#### Supplemental Table 2. Healthy donor characteristics

| Donor | Bat ch | 10x chemistry | Age (years) | Sex | Race/ethnicity (self-reported) | Race/ethnicity group |
|---|---|---|---|---|---|---|
| HDonor1 | 1 | Chromium Single Cell 3′ v3.1 | 51 | M | Caucasian | White |
| HDonor2 | 1 | Chromium Single Cell 3′ v3.1 | 34 | F | Caucasian | White |
| HDonor3 | 1 | Chromium Single Cell 3′ v3.1 | 33 | M | Asian | Asian |
| HDonor4 | 1 | Chromium Single Cell 3′ v3.1 | 41 | M | White | White |
| HDonor5 | 1 | Chromium Single Cell 3′ v3.1 | 32 | M | Asian | Asian |
| HDonor6 | 1 | Chromium Single Cell 3′ v3.1 | 23 | F | Asian | Asian |
| HDonor7 | 1 | Chromium Single Cell 3′ v3.1 | 34 | F | White | White |
| HDonor8 | 1 | Chromium Single Cell 3′ v3.1 | 36 | M | Caucasian | White |
| HDonor9 | 1 | Chromium Single Cell 3′ v3.1 | 26 | M | Asian | Asian |
| HDonor10 | 2 | Chromium Single Cell 3′ v3.1 | 26 | F | Asian | Asian |
| HDonor11 | 2 | Chromium Single Cell 3′ v3.1 | 25 | F | Asian | Asian |
| HDonor12 | 2 | Chromium Single Cell 3′ v3.1 | 30 | F | White/Latina | Hispanic/Latino |
| HDonor13 | 2 | Chromium Single Cell 3′ v3.1 | 31 | M | Asian | Asian |
| HDonor14 | 2 | Chromium Single Cell 3′ v3.1 | 47 | M | White | White |
| HDonor15 | 2 | Chromium Single Cell 3′ v3.1 | 32 | M | Asian | Asian |
| HDonor16 | 2 | Chromium Single Cell 3′ v3.1 | 32 | M | Asian | Asian |
| HDonor17 | 2 | Chromium Single Cell 3′ v3.1 | 26 | F | Hispanic | Hispanic/Latino |
| HDonor18 | 2 | Chromium Single Cell 3′ v3.1 | 30 | M | White | White |
| HDonor19 | 2 | Chromium Single Cell 3′ v3.1 | 34 | F | White | White |
| HDonor20 | 2 | Chromium Single Cell 3′ v3.1 | 37 | F | Asian | Asian |
| HDonor21 | 3 | Chromium Single Cell 3′ v3.1 | 29 | F | Middle Eastern | Middle Eastern/North African |
| HDonor22 | 3 | Chromium Single Cell 3′ v3.1 | 50 | F | Southern European and North African | White |
| HDonor23 | 4 | GEM-X Single Cell 3′ v4 | 53 | M | White | White |
| HDonor24 | 4 | GEM-X Single Cell 3′ v4 | 47 | M | Caucasian | White |
| HDonor25 | 4 | GEM-X Single Cell 3′ v4 | 54 | F | White | White |
| HDonor26 | 4 | GEM-X Single Cell 3′ v4 | 52 | F | Romanian | White |
| HDonor27 | 4 | GEM-X Single Cell 3′ v4 | 45 | M | White | White |
| HDonor28 | 4 | GEM-X Single Cell 3′ v4 | 55 | F | White | White |
| HDonor29 | 4 | GEM-X Single Cell 3′ v4 | 39 | M | White | White |
| HDonor30 | 4 | GEM-X Single Cell 3′ v4 | 46 | M | Middle Eastern | Middle Eastern/North African |
| HDonor31 | 4 | GEM-X Single Cell 3′ v4 | 59 | F | Middle Eastern | Middle Eastern/North African |
| HDonor32 | 4 | GEM-X Single Cell 3′ v4 | 47 | M | Egyptian | Middle Eastern/North African |
| HDonor33 | 4 | GEM-X Single Cell 3′ v4 | 44 | F | Filipino/Asian | Asian |
| HDonor34 | 4 | GEM-X Single Cell 3′ v4 | 47 | F | Irish | White |

#### Supplemental Table 3. Sepsis pilot cohort: clinical characteristics

| Participant | Group | Age (years) | Sex | Race/ethnicity | Main reason for ICU admission | In purified analyses |
|---|---|---|---|---|---|---|
| H_1 | Control | 69 | M | White | Severe asthma | Yes |
| H_2 | Control | 85 | F | White | Airway obstruction due to malignancy | Yes |
| H_3 | Control | 47 | M | White | — | No |
| H_4 | Control | 34 | F | White | — | Yes |
| S_1 | Sepsis | 72 | M | Other | Septic shock, urinary source (E. coli bacteremia) | Yes |
| S_2 | Sepsis | 57 | M | Asian | Septic shock; suspected invasive fungal sinusitis | Yes |
| S_3 | Sepsis | 60 | M | Other | Shock (septic vs neurogenic) with flaccid paralysis | Yes |
| S_4 | Sepsis | 83 | F | Asian | Septic shock with lactic acidosis and acute kidney injury | No |

### Supplemental Table 4. Software, versions and key parameters

| Software | Version | Used for | Key parameters |
|---|---|---|---|
| Cell Ranger | 8.0.1 | Read alignment and UMI counting, all Siphon-seq libraries | cellranger count, default settings; reference GRCh38-2024-A (10x refdata-gex-GRCh38-2024-A) |
| Python | 3.13 | Single-cell processing and classification (human) | conda env concord_2 |
| scanpy | 1.11.1 | Normalization, HVGs, PCA, neighbors, UMAP, Leiden, marker tests | normalize_total (median), log1p, 2,000 HVGs (batch_key = donor, i.e. the donor library), 50 PCs, Leiden (igraph, resolution 1, 2 iterations), Wilcoxon rank_genes_groups |
| anndata | 0.12.1 | Data objects | |
| CONCORD (concord_sc) | 0.9.3 | Partition classifier (platelet / RBC / WBC) | 1,387 input genes; 50,000 GEMs per class; domain key = donor library; classifier head; train_frac 0.9; 10 epochs; latent 32; encoder [128]; batch 64; lr 0.01; augmentation mask 0.5; seed 0 |
| PyTorch | 2.6.0 | CONCORD backend (CPU) | |
| umap-learn | 0.5.7 | UMAP of the CONCORD latent | n_neighbors 15, min_dist 0.1, Euclidean |
| python-igraph / leidenalg | 0.11.8 / 0.10.2 | Leiden clustering | |
| gseapy | 1.1.3 | Preranked GSEA (sepsis pilot, GSE252275) | GO Biological Process 2023, Reactome 2022; min 15 / max 500 genes (predefined panel: min 8); 10,000 permutations; seed 42 |
| pyDESeq2 | 0.5.4 | Sepsis vs control differential expression (sepsis pilot) | design ~group; Wald test (alpha 0.1, Cooks refit); ranking by Wald statistic |
| R | 4.4.1 | Dispersion tiering, MuSiC, edgeR | |
| edgeR / limma | 4.2.1 / 3.60.4 | TMM normalization, CPM, dispersion; mouse exact test | TMM |
| MuSiC | 1.0.0 | Cell-type RNA fractions (all bulk and pseudobulk libraries) | music_prop with default arguments against the six-type reference; output converted to RNA share |
| babelgene | 22.9 | Human–mouse 1:1 orthologs (mouse experiment) | |
| scipy / numpy / pandas | 1.15.2 / 2.1.3 / 2.3.3 | Statistics and tables (figure scripts) | two-sided tests throughout |
| matplotlib | 3.10 | Figures | |

### Supplemental Table 5. Public platelet RNA-seq projects

| Project | BioProject / ENA | GEO | Reference | Condition | Anticoagulant | Platelet isolation | Leukocyte removal | Library | Samples (n) | Healthy samples (n) |
|---|---|---|---|---|---|---|---|---|---|---|
| P01 | PRJNA521955 | GSE126446 | Bongiovanni 2019[30] | Healthy; immature vs mature platelets | NR | FACS sort on thiazole orange (top 15% vs bottom 30%) | FACS sort | SMART-Seq Stranded, total RNA; SE 75 | 8 | 8 |
| P02 | PRJEB33335 | — | Hille 2020[31] | Healthy; reticulated vs non-reticulated platelets | NR | Flow sort on SYTO 13 | Flow sort | Long RNA-seq, kit NR; PE 76 | 12 | 12 |
| P03 | PRJNA722042 | — | Bao 2021[32] | Acute myeloid leukemia | NR | As Eicher 2016 (cited) | None reported | Oligo(dT)-primed, low input (abstract: TruSeq Stranded mRNA); PE | 13 | 0 |
| P05 | PRJNA595551 | — | Frey 2020[33] | Advanced heart failure, before and after LVAD | ACD | PRP (150 × g, 20 min), then leukocyte depletion | Leukocyte-depleted (< 3 per 10^7 platelets) | TruSeq, poly(A); SE 50 | 34 | 0 |
| P06 | PRJNA274751 | GSE65705 | Eicher 2016[34] | Acute myocardial infarction (STEMI, NSTEMI) | Citrate | CPT tube (1,700 × g); platelets from supernatant of 288 × g spin | | NuGEN Ovation RNA-Seq v2, total RNA; PE 75 | 32 | 0 |
| P07 | PRJEB35689 | — | Hille 2020[31] | Healthy, chronic coronary syndrome, STEMI; reticulated vs non-reticulated platelets | NR | Flow sort on SYTO 13 | Flow sort | Long RNA-seq, kit NR; PE 75 | 48 | 16 |
| P08 | PRJNA664615 | — | Xu C 2022[35] | Immune thrombocytopenia | EDTA | Differential centrifugation (PRP, then platelet pellet) | | Oligo(dT), as Best 2017; PE 150 | 59 | 0 |
| P09 | PRJNA421730 | GSE107868 | Best 2019[36] | Lower-grade glioma, healthy | EDTA | thromboSeq (PRP 120 × g, pellet 360 × g) | | SMARTer Ultra Low v3, TruSeq; SE 100 | 80 | 41 |
| P10 | PRJNA531691 | — | Frey 2020[33] | Healthy, sampled longitudinally | Citrate or ACD | CD45 microbead depletion | CD45 microbeads | KAPA or TruSeq, poly(A); SE 50 | 88 | 88 |
| P11 | PRJNA281708 | GSE68086 | Best 2015[37] | Six cancer types, healthy | EDTA | thromboSeq | | SMARTer Ultra Low v1, TruSeq; SE 100 | 266 | 55 |
| P12 | PRJNA737596 | — | Xu L 2022[38] | Colorectal cancer, intestinal disease, healthy | EDTA | thromboSeq (cited) | | SMARTer, Ovation SoLo; PE 100 | 322 | 21 |
| P13 | PRJNA353588 | GSE89843 | Best 2017[39] | Non-small-cell lung cancer, non-cancer controls | EDTA | thromboSeq | | SMARTer Ultra Low v3, TruSeq Nano; SE 100 | 645 | 228 |
| P14 | PRJNA659491 | GSE156902 | Sol 2020[40] | Glioblastoma, brain metastases, multiple sclerosis, controls | EDTA | thromboSeq | | SMARTer Ultra Low v3, TruSeq Nano; SE 100 | 765 | 335 |
| P15 | PRJNA761450 | GSE183635 | In 't Veld 2022[39,41] | 18 cancer types, controls | EDTA | thromboSeq | | SMARTer Ultra Low v3, TruSeq Nano; SE 100 | 1317 | 34 |

### Supplemental Figures

#### Supplemental Figure 1. Public-data deconvolution reference and validation

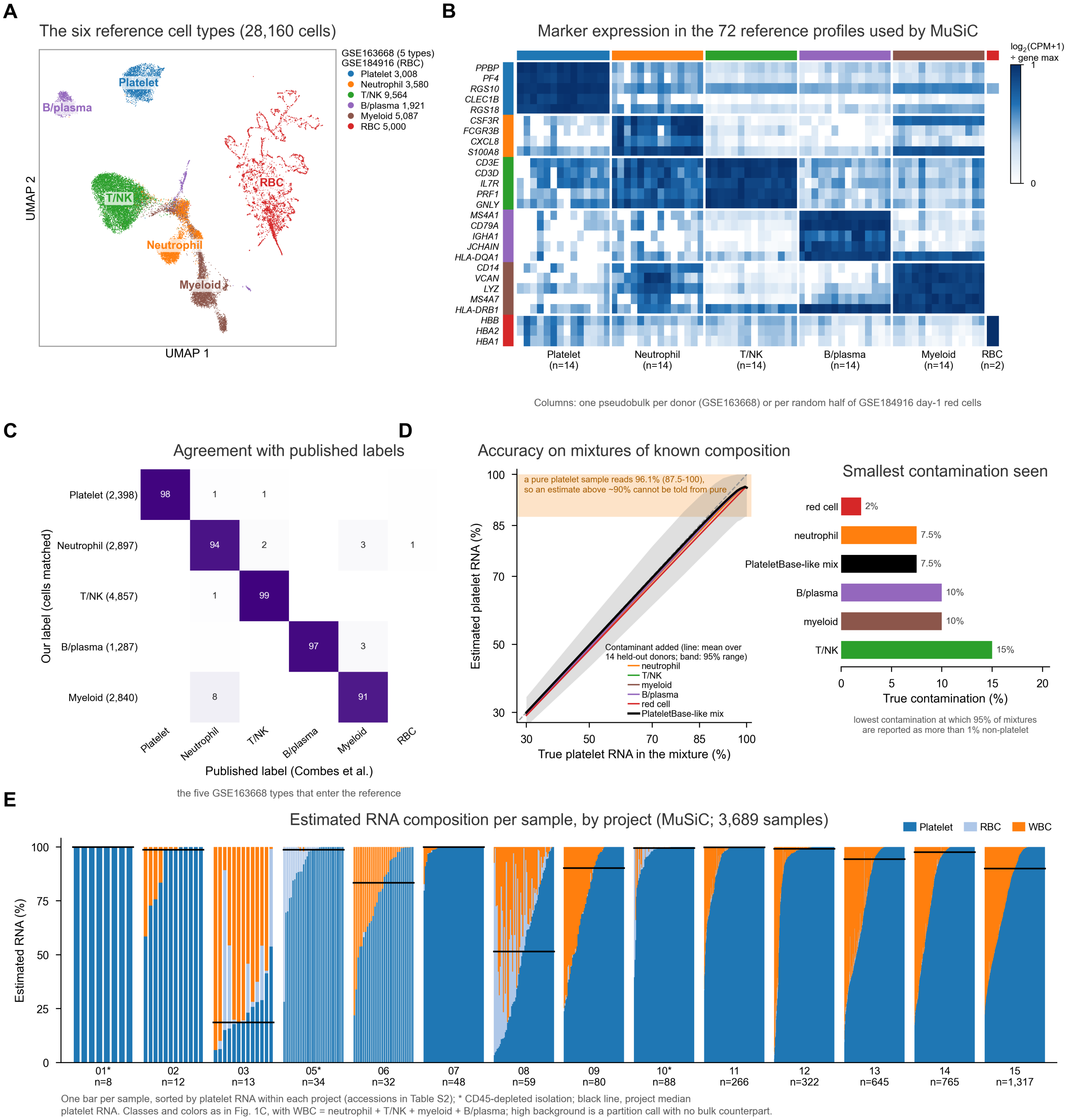


**Figure S1. The single-cell reference used for MuSiC deconvolution, its accuracy, and per-sample platelet RNA in the public projects.**

The reference combines platelets and immune cells from healthy-donor whole-blood scRNA-seq (Combes et al., Nature 2021; GSE163668) with erythrocytes from a separate red-cell dataset (Jain et al. 2022; GSE184916, day-1 sample).

**(A)** Embedding of the 28,160 cells that make up the reference: the five cell types taken from the 14 healthy-donor libraries of GSE163668, after independent reproduction of the authors' coarse annotation (SCTransform, Harmony across donors, Louvain at resolution 0.6, and marker module scores with manual review), together with the 5,000 day-1 red cells of GSE184916. Cells are colored by cell type and the legend gives the number of each. Red cells were taken from a second dataset because GSE163668 lysed red cells before capture: its red-cell-labeled cluster proved to be mostly neutrophils carrying ambient hemoglobin RNA, and those cells, along with an ambiguous and a low-complexity cluster, are not part of the reference and are not shown. The two datasets were embedded together without batch correction, so the red cells sit apart both because they are a different cell type and because they come from a different experiment; the embedding is for display only.

**(B)** Expression of the authors' marker panels in the 72 pseudobulk profiles that form the MuSiC reference (10,132 genes). There are 14 donor profiles each for platelet, neutrophil, T/NK, B/plasma and myeloid cells (GSE163668), and 2 red-cell profiles from random halves of the GSE184916 day-1 sample. Values are log2(CPM + 1), with each gene scaled to its maximum. Three panel genes absent from the reference gene space (G0S2, HBD, HBM) are not shown.

**(C)** Agreement of our labels with the published per-cell labels for the same libraries, for the five GSE163668 cell types that enter the reference (21,292 cells matched by barcode, of which 14,279 carry one of these five labels). Each row gives the percentage of cells with our label that carry each published label; agreement is 91–99% for every type.

**(D)** Accuracy of MuSiC platelet RNA estimates on mixtures of known composition. *Left:* the reference was rebuilt four times, each time from a subset of donors, and platelet profiles from the held-out donors were mixed in count space with held-out contaminant profiles, either one cell type at a time (neutrophil, T/NK, myeloid, B/plasma or red cell) or a PlateletBase-like mixture (neutrophil 43%, T/NK 21%, red cell 20%, myeloid 12%, B/plasma 3%), at true platelet RNA shares of 30–100%, downsampled to 3.8 million counts, the median PlateletBase library; 14 donors × 5 draws per point. Each mixture was then fitted with the same call used for every sample in this study. Lines, mean estimate over the held-out donors; gray band, 95% range for the PlateletBase-like mixture; dashed line, perfect agreement. Estimates track the true value with a root-mean-square error of 0.05–0.06 between 85% and 100% true platelet RNA and 0.02–0.03 at 30%, and the error is set by which donor is held out rather than by sequencing depth (it changed by ≤ 0.003 at 0.8 million counts). The orange band shows the estimates for pure platelet mixtures (mean 96.1%, 95% range 87.5–100%), so an estimate above about 90% cannot be distinguished from pure platelet RNA, while one below about 85% indicates contamination. *Right:* detection limit for each contaminant, defined as the lowest true contamination at which at least 95% of mixtures are estimated to hold more than 1% non-platelet RNA. Red cells are detected from 2% because their transcriptome is almost entirely hemoglobin, and T/NK cells from 15%.

**(E)** Estimated RNA composition of every sample in the 14 PlateletBase projects (accessions in Supplemental Table 5): one stacked bar per sample, sorted by platelet RNA within each project. Leukocyte (WBC) is the sum of the neutrophil, T/NK, myeloid and B/plasma shares; high background is a partition call made on single GEMs and has no counterpart in a bulk library. The black line is the project median platelet RNA and asterisks mark CD45-depleted isolation. Composition varies widely within projects as well as between them: project 08 is bimodal and project 03 runs from below 10% to about 50% platelet RNA, spreads far larger than the error in (D). The main contaminant also differs by project: leukocyte RNA in projects 09 and 13–15, and red-cell RNA in project 05 and parts of 03 and 08. Project 04 (n = 20) was excluded for insufficient depth, and two project-06 libraries with fewer than 1,000 reads were not fitted. 9,738–9,748 reference genes were shared with each project (7,729 for project 15). Estimates of exactly 100% (1,186 samples) are at the non-negativity boundary and are censored, not measured.

### Supplemental Figure 2. Partition classification

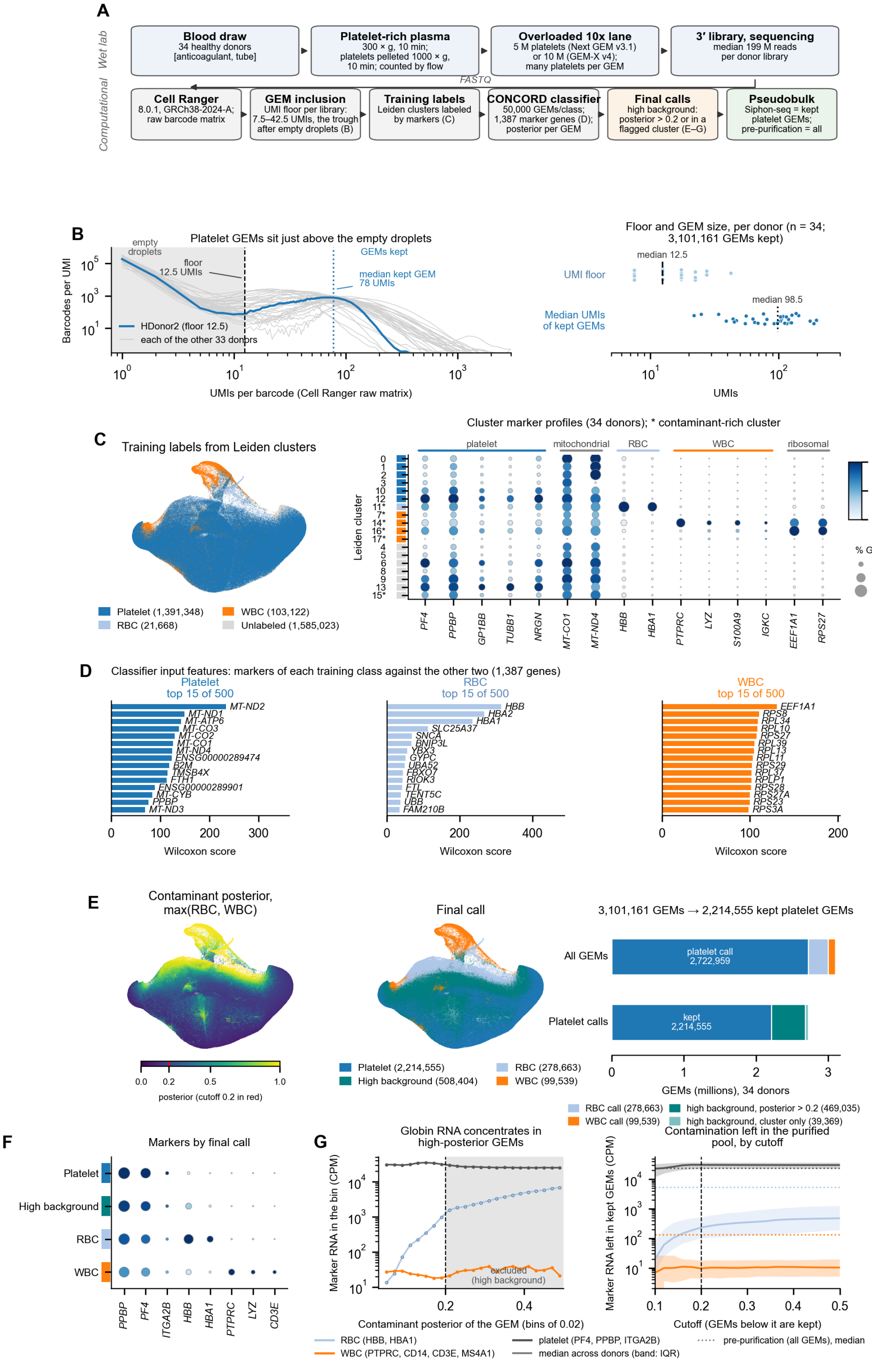


**Figure S2. Siphon-seq partition classification, step by step.** Each panel shows one step of the workflow in (A), for the 34 healthy-donor libraries. Panels C and E are drawn on the same two-dimensional CONCORD UMAP; plotted points are a random 1,000,000 of the 3,101,161 GEMs.

**(A)** The Siphon-seq workflow, wet lab above and computational below; each computational step names the panel that shows it. Parameters are those recorded in the processing notebooks (Supplemental Methods, Partition classification).

**(B)** GEM inclusion. Number of barcodes at each UMI count in the Cell Ranger raw matrices, for one representative donor (blue; HDonor2, the library at the median floor) and each of the other 33 donors (gray). Platelet GEMs carry far less RNA than nucleated cells, and form a shoulder immediately above the empty-droplet peak rather than a well-separated knee, which is why Cell Ranger's own cell calling was not used. For each library the inclusion threshold was set at the trough between the two, the first 5-UMI bin in which the slope of the histogram turns positive (dashed line; 13 UMIs for this donor). The dotted line is the median UMI count of the GEMs that library keeps (78), about six times the threshold. Right: both values for every donor, the floor (median 12.5, range 7.5–42.5) and the median UMIs of the kept GEMs (median 98.5, range 22–197). In total, 3,101,161 GEMs were retained.

**(C)** Training labels. Left: Leiden clusters of the combined object, colored by the label used to train the classifier; clusters that did not carry unambiguous markers of one lineage were left unlabeled and were not used for training. Right: each cluster's marker profile, as the percentage of its GEMs with any count of a gene (dot size) and the cluster's mean expression scaled to the highest cluster for that gene (color). Clusters 11 and 7, 14, 16, 17 carry erythrocyte and leukocyte markers respectively; the platelet-labeled clusters differ mainly in mitochondrial content. Asterisks mark the clusters later flagged as contaminant-rich.

**(D)** Classifier input features. The 15 highest-scoring marker genes of each training class against the other two, by Wilcoxon rank-sum test on a balanced draw of 50,000 GEMs per class. The classifier's input is the union of the top 500 per class. Erythrocyte GEMs are defined by hemoglobin and leukocyte GEMs largely by ribosomal-protein transcripts, while the platelet classes' top genes are mitochondrial, reflecting the composition of the platelet transcriptome rather than a marker chosen by hand.

**(E)** Final calls. Left: the contaminant posterior of each GEM, the larger of its erythrocyte and leukocyte posteriors; the 0.2 cutoff is marked in red on the color bar. Center: the final call per GEM. Right: how the GEMs divide at each step. A platelet-called GEM was reclassified as high background, and excluded, if its contaminant posterior exceeded 0.2 or it lay in a cluster in which more than half of GEMs did.

**(F)** Mean expression and percentage of GEMs detected for lineage markers, by final call. High- background GEMs carry platelet markers together with the contaminant markers, which is what distinguishes them from the erythrocyte and leukocyte calls.

**(G)** Choice of the 0.2 cutoff. Left: marker RNA, as counts per million, in GEMs grouped by contaminant posterior (0.02 bins, 34 donors pooled, GEMs in contaminant-rich clusters excluded). Globin RNA rises steeply with the posterior while platelet marker RNA does not. Right: marker RNA remaining in the retained GEMs at each candidate cutoff (line, median across the 34 donors; band, interquartile range; dotted line, the pre-purification library). Leukocyte marker RNA is flat above about 0.14, so the cutoff mainly governs how much globin RNA is removed.

#### Supplemental Figure 3. RNA share versus GEM share

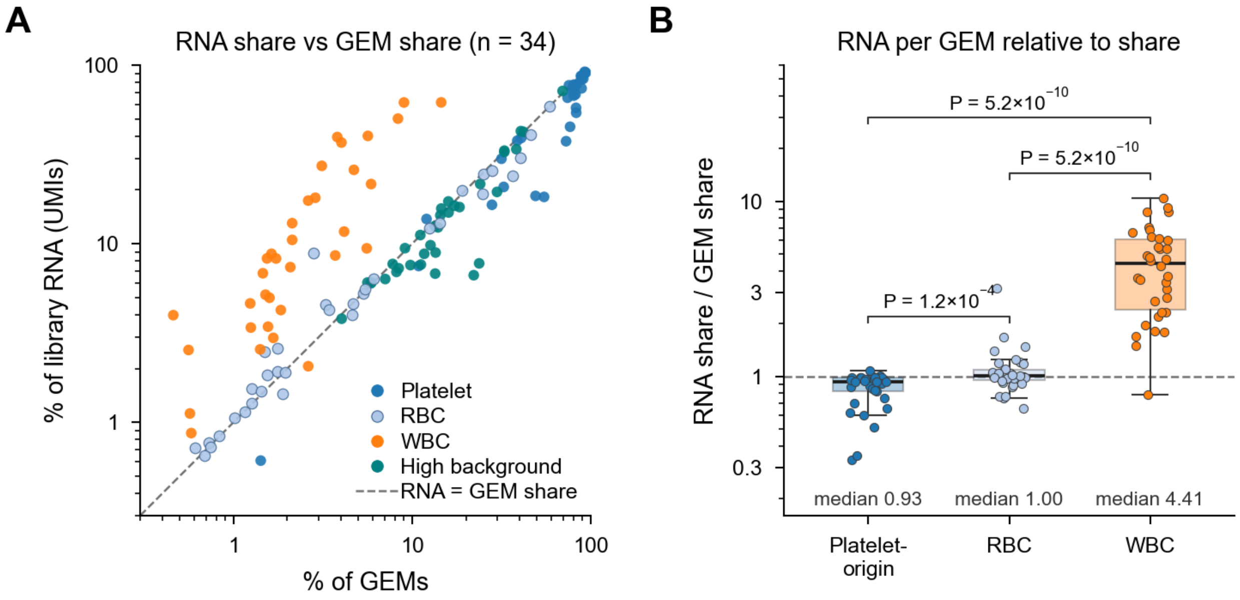


**Leukocyte-containing GEMs contribute more RNA than their number predicts.** Siphon-seq partitions from 34 healthy donors (10x 3′) were classified as platelet, erythrocyte (RBC), leukocyte (WBC) or high background. For each donor, a class's GEM share is its fraction of all GEMs and its RNA share is its fraction of all UMIs in the pseudobulk library. **(A)** RNA share vs GEM share per donor and class (n = 34 donors; log scales). Dashed line, equal shares. WBC GEMs lie above the line. **(B)** RNA-to-GEM share ratio per donor, with platelet and high-background GEMs pooled as platelet-origin. A ratio of 1 (dashed line) means a class contributes RNA in proportion to its GEM count. Medians: platelet-origin 0.93, RBC 1.00, WBC 4.41. Boxes show the median and IQR, and whiskers extend to the range. Two-sided paired Wilcoxon signed-rank tests with Benjamini–Hochberg adjustment over three comparisons (n = 34). Counting RNA rather than GEMs therefore lowers the apparent platelet content of a preparation: the platelet share falls from a median of 80.3% of GEMs to 70.7% of library RNA, and is lower in RNA in 32 of 34 donors (two-sided paired Wilcoxon signed-rank test, $P = 8.1 \times 10^{-9}$). The donors that lose most are those with the most leukocyte GEMs, up to 36 percentage points in the donor whose 9.0% leukocyte GEMs carry 61.8% of the library (54.8% of GEMs platelet, 18.3% of RNA). Per-donor values are in Supplemental Data 2.

### Supplemental Figure 4. Siphon-seq in mouse blood

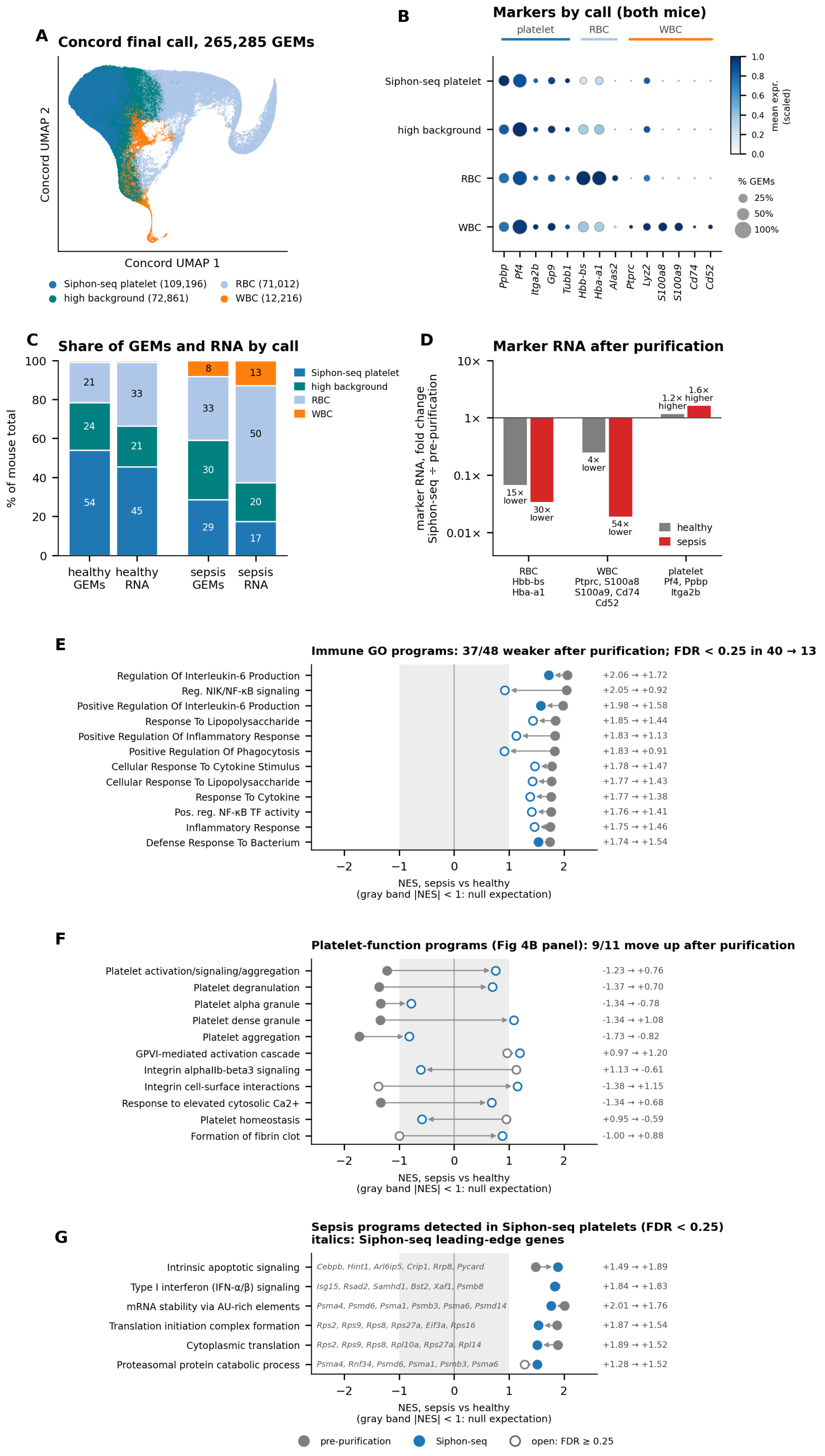


**Figure S4. Siphon-seq purifies platelet GEMs from mouse blood and removes contaminant-driven signal from a sepsis comparison.** Blood from three mice three days after cecal-slurry injection (sepsis) and three vehicle-injected controls (healthy) was pooled by condition and processed by Siphon-seq as one library per condition; GEMs were classified with a CONCORD classifier trained on marker-labeled Leiden clusters, exactly as for the human donors. **(A) CONCORD UMAP** of all 265,285 GEMs from both libraries, colored by final call. Platelet-predicted GEMs with a contaminant (RBC or WBC) posterior > 0.2, or in Leiden clusters in which > 50% of GEMs exceeded that posterior, are called high background and excluded; the remaining platelet GEMs form the

Siphon-seq profile. **(B)** Marker expression by call (both libraries combined). Dot size, % of GEMs with ≥ 1 UMI; color, mean log-normalized expression scaled to each gene's maximum. **(C)** Share of GEMs and of RNA (UMIs) in each call, per library. **(D)** Fold change in marker RNA after purification (CPM in the Siphon-seq profile ÷ CPM in the pre-purification profile of all GEMs), per library. RBC markers Hbb-bs, Hba-a1; WBC markers Ptprc, S100a8, S100a9, Cd74, Cd52 (the WBC genes of panel B, excluding Lyz2, which is abundant in mouse platelet GEMs); platelet markers Pf4, Ppbp, Itga2b. **(E)** Normalized enrichment scores (NES) for sepsis vs healthy in the pre-purification (gray) and Siphon-seq (blue) profiles for immune GO Biological Process terms. Rows are the 12 immune terms with the highest pre-purification NES among those at FDR < 0.25 pre-purification; the title gives counts over all 48 immune terms tested. **(F)** The same comparison for the platelet-function and procoagulant programs of the predefined platelet panel. **(G)** Sepsis programs detected in the Siphon-seq profile: every program of the predefined platelet panel with FDR < 0.25 in Siphon-seq, plus Reactome type I interferon (IFN-α/β) signaling. Italics, Siphon-seq leading-edge genes. In E–G, filled points have FDR < 0.25 and the gray band marks |NES| < 1, the null expectation. Genes were ranked by log2 fold change (sepsis/healthy) after DESeq2 median-of-ratios scaling, restricted to genes with ≥ 10 UMIs in both libraries and mapped to human 1:1 orthologs; preranked GSEA used GO Biological Process 2023 (E), the predefined panel (F, G) or Reactome 2022 (G, interferon), 10,000 permutations. With one pooled library per condition, all comparisons are descriptive.

#### Supplemental Figure 5. Contextual genes outside the core

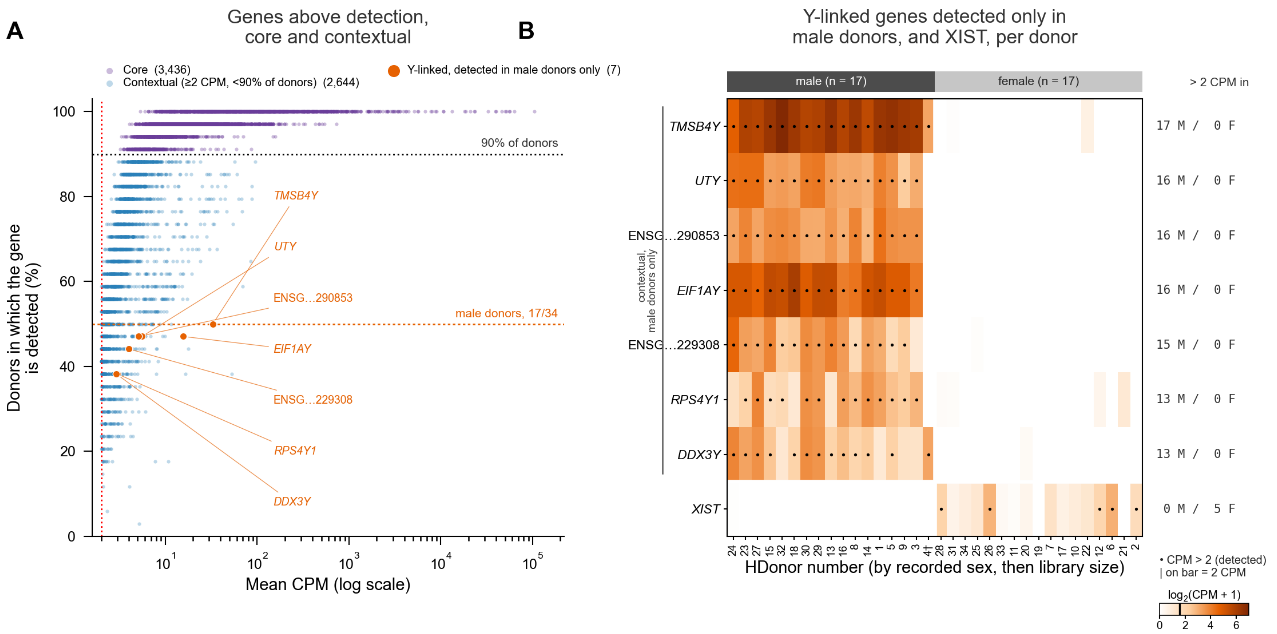


**Figure S5. Genes above the detection threshold that fall outside the core.**

(A) Mean TMM-CPM across 34 donors versus the percentage of donors in which each gene exceeds 2 CPM, for the 6,080 genes with mean CPM ≥ 2 (without the 32,526 genes below detection). Purple, core (3,436). Blue, contextual (2,644: mean ≥ 2 CPM but detected in < 90% of donors). Orange, labeled, the 7 Y-linked genes detected only in male donors: *TMSB4Y*, *UTY*, *EIF1AY*, *RPS4Y1*, *DDX3Y*, and the lncRNAs ENSG00000290853 and ENSG00000229308 (GENCODE v47; pseudoautosomal genes excluded). Dotted lines, the two core criteria (mean ≥ 2 CPM; detected in ≥ 90% of donors). Orange dotted line, the share of male donors (17 of 34, 50%). Contextual genes are spread across the whole range of donor prevalence rather than sitting just below the core threshold, and are almost all autosomal: of the 2,644, 2,498 are autosomal, 88 X-linked, 10 Y-linked, 5 pseudoautosomal and 1 maps to two autosomes, while 42 gene names in the count matrix are not in GENCODE v47.

(B) Per-donor TMM-CPM ($\log_2(\text{CPM} + 1)$) of the 7 male-only Y-linked genes and of *XIST*. Donors are grouped by recorded sex and ordered by library size within sex; dots, CPM > 2. Right, number of male (M) and female (F) donors with CPM > 2; the black mark on the color bar is the 2 CPM detection threshold. Each gene exceeds 2 CPM in 13–17 of the 17 male donors and in no female donor, so it is detected in at most 50% of donors and cannot reach the 90% core criterion. *XIST* is present at low levels in female donors (mean 1.8 CPM) and absent in male donors, but exceeds the 2 CPM detection threshold in only 5 of 17 female donors (mean across all donors 0.88 CPM, below detection). Donor sex is as recorded in the donor metadata. Expression of six male-specific Y genes agrees with it in 33 of 34 donors. †HDonor4 has a male-level rate, but from only 2 reads.

### Supplemental Figure 6. Megakaryocyte expectation models

##### Core membership model (Figure 2C, 2E, 2F)

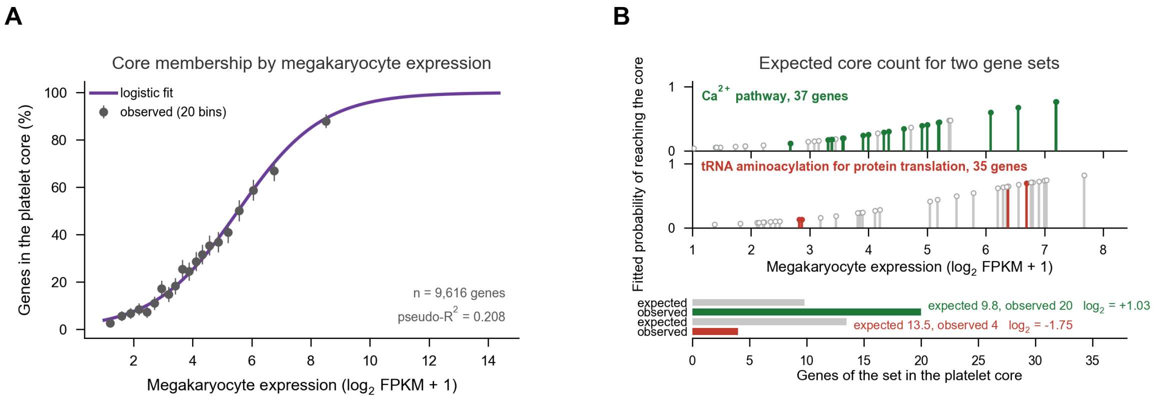


##### Transcript amount model (Figure 2G; enrichment axis of Figure 2H)

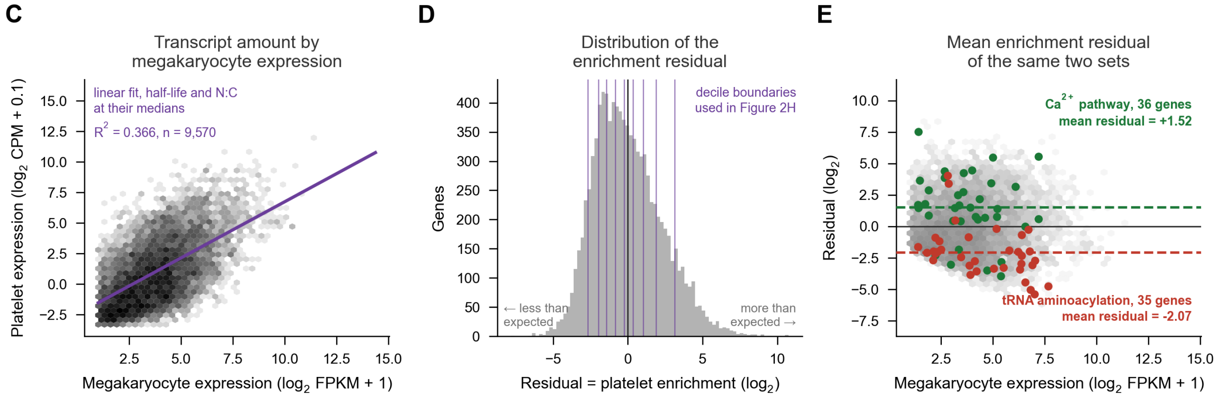


##### Protein detection model (expected detection in Figure 2H)

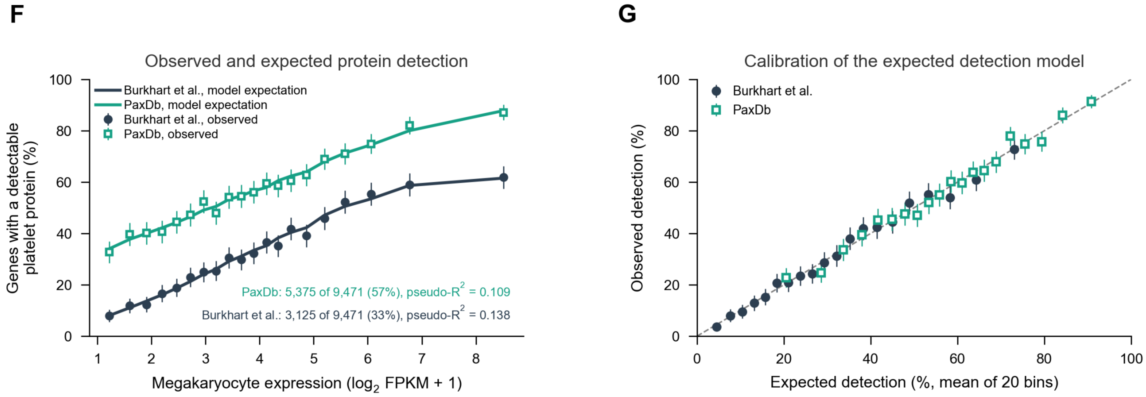

**Figure S6. Megakaryocyte expectation models.** Each model predicts a platelet readout from megakaryocyte expression (BLUEPRINT), with mRNA half-life and nuclear:cytoplasmic (N:C) ratio added in (C) and (F); gene sets are scored against these predictions.

**(A)** Percentage of genes that reach the platelet core, by megakaryocyte expression (BLUEPRINT, $\log_2$ FPKM + 1). Points, 20 equal-sized bins with 95% confidence intervals; line, the fitted logistic regression (n = 9,616 genes; McFadden pseudo-$R^2$ = 0.208). This fit supplies the expected core membership against which gene sets are scored.

**(B)** Construction of the expectation for two gene sets of almost the same size that fall on opposite sides of it: the $Ca^{2+}$ pathway (Reactome, 37 genes; green) and tRNA aminoacylation for protein translation (GO, 35 genes; red). *Top:* each member gene is a stick placed at its megakaryocyte expression, with height equal to its fitted probability of reaching the core from (A); colored sticks are the genes that are in the core, gray sticks the genes that are not. *Bottom:* the expected count for each set (the sum of those probabilities, that is, the summed height of its sticks) against the number observed. The $Ca^{2+}$ pathway contributes 20 core genes against 9.8 expected ($\log_2$ observed/expected = +1.03), and tRNA aminoacylation 4 against 13.5 expected ($\log_2$ = −1.75). The two sets are matched for size and span a similar range of megakaryocyte expression, so their opposite departures are not explained by megakaryocyte expression level.

**(C)** Platelet transcript amount ($\log_2$ CPM + 0.1) against megakaryocyte expression, for the 9,570 genes with any platelet reads. Line, the linear fit with mRNA half-life and N:C ratio held at their medians ($R^2$ = 0.366 for the full three-covariate model). The residual of this fit is defined as platelet enrichment: one unit is a twofold departure from the amount the three covariates predict.

**(D)** Distribution of that residual across genes. Purple lines, the decile boundaries used in the protein-detection analysis; the black line is zero, where a gene is present in exactly the amount predicted.

**(E)** The same two gene sets on the residual scale, against all genes (gray). The $Ca^{2+}$ pathway sits above the line of no enrichment (mean residual +1.52, a 2.9-fold excess of transcript over the amount predicted; $q = 3.6 \times 10^{-4}$) and tRNA aminoacylation below it (−2.07, a 4.2-fold shortfall; $q = 1.4 \times 10^{-6}$). The sets depart in the same directions here as in (B), so the two statistics (whether a gene reaches the core at all, and how much of it is present) agree for these sets, although they are computed differently. The $Ca^{2+}$ pathway has 36 genes here against 37 in (B), because the transcript-amount model is fitted on the 9,570 genes with any platelet reads rather than the 9,616 of the core model.

**(F)** Percentage of genes whose protein is detected in the platelet, by megakaryocyte expression, for each proteome: Burkhart et al. (3,125 of 9,471 genes detected, 33%) and PaxDb (5,375, 57%). Points, 20 equal-sized bins with 95% confidence intervals; lines, the mean fitted probability of the genes in the same bins under a logistic model of detection on a B-spline in megakaryocyte expression plus half-life and N:C ratio (McFadden pseudo-$R^2$ 0.138 and 0.109). This model contains no platelet measurement, so it supplies the expected detection independently of platelet enrichment.

**(G)** Calibration of that model: observed detection against mean expected detection, in 20 bins of predicted probability, with the line of equality. Expected detection tracks observed detection across the whole range.

#### Supplemental Figure 7. Purity gradient across datasets

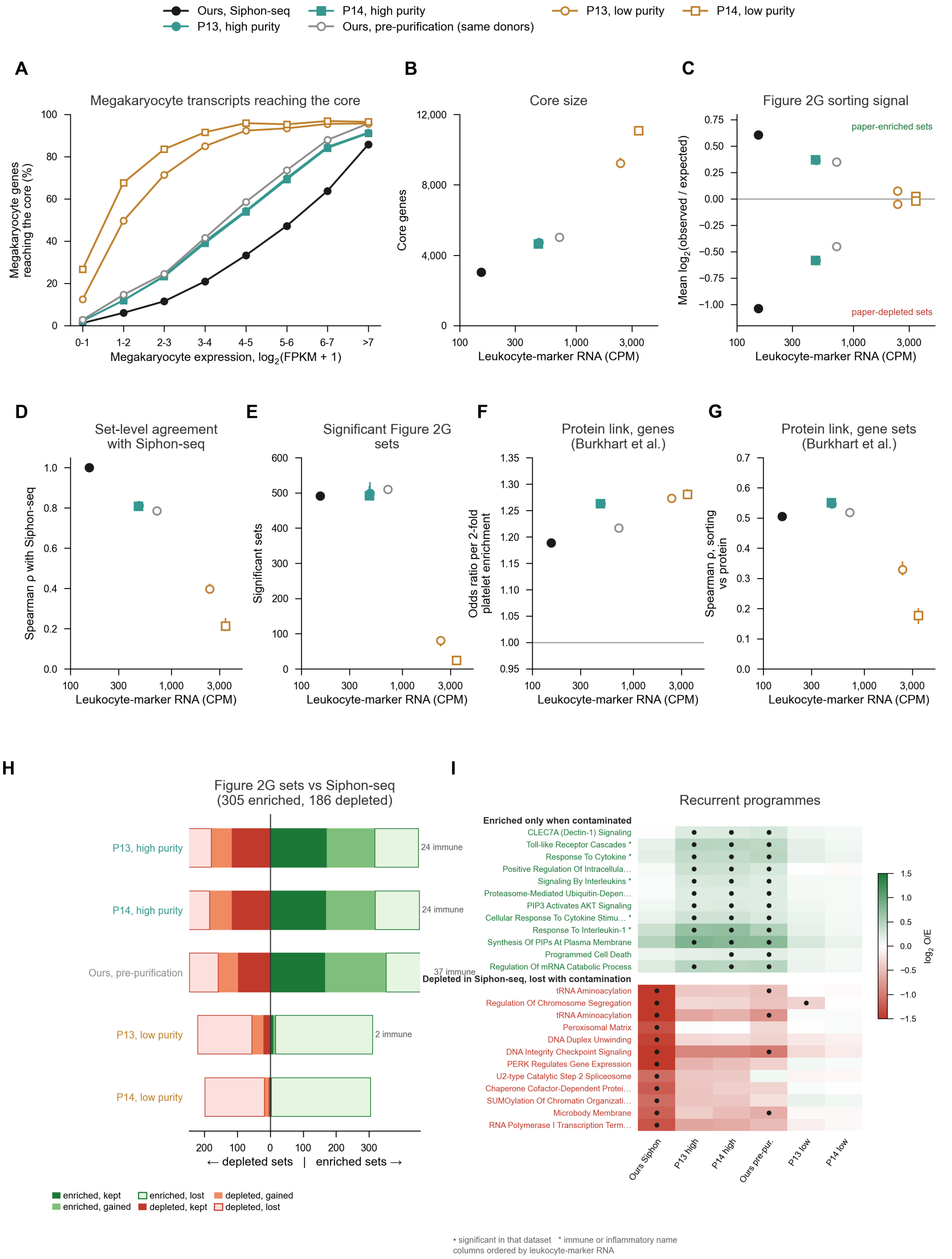


**Figure S7. Gene-set sorting readouts across platelet datasets ordered by leukocyte RNA content.** Six platelet transcriptomes are compared: Siphon-seq (34 donors), the same 34 libraries before purification, and the high-purity and low-purity halves of two public PlateletBase projects of healthy donors split by MuSiC platelet proportion (P13, PRJNA353588, 110 + 110 samples; P14, PRJNA659491, 160 + 160 samples). All datasets are scored on the 18,201 gene symbols quantified in every dataset, using the core rule of the main text (mean CPM ≥ 2 and CPM > 2 in ≥ 90% of samples, after TMM normalization), so set counts here differ slightly from those on the full universe. Values for the public halves are medians of 20 random draws of 34 samples, with bars giving the interquartile range; the two arms of this study are single values at n = 34. Leukocyte-marker RNA, the x-axis of (B) to (G), is the median CPM of leukocyte markers: 153 CPM in Siphon-seq, 478 and 472 in the P13 and P14 high-purity halves, 717 before purification, and 2,387 and 3,420 in the low-purity halves.

**(A)** Percentage of megakaryocyte-quantified genes reaching the core, by megakaryocyte expression (BLUEPRINT, $\log_2$ FPKM + 1, eight bins). Of genes expressed at 3 to 4, 21.0% reach the Siphon-seq core, against 39 to 42% in the high-purity halves and before purification, and 85 to 92% in the low-purity halves.

**(B)** Core size on the common universe: 3,040 genes in Siphon-seq, 4,719 and 4,653 in the high-purity halves, 5,024 before purification, and 9,225 and 11,079 in the low-purity halves.

**(C)** Mean $\log_2$(observed/expected) core membership of the enriched and depleted sets called in Siphon-seq on the full universe, with the megakaryocyte model refitted in each dataset. Both shifts collapse toward zero with contamination: enriched sets +0.61 in Siphon-seq, +0.37 in both high-purity halves, +0.35 before purification, +0.08 and +0.03 in the low-purity halves; depleted sets −1.04, −0.59 and −0.58, −0.45, −0.05 and −0.02.

**(D)** Spearman ρ between each dataset's set-level $\log_2$ observed/expected and that of Siphon-seq: 0.81 in both high-purity halves, 0.79 before purification, 0.40 and 0.21 in the low-purity halves.

**(E)** Number of significant sets (Benjamini-Hochberg $q < 0.05$): 491 in Siphon-seq, 499 and 492 in the high-purity halves, 510 before purification, but 80.5 and 24.5 in the low-purity halves. Moderate contamination changes which sets are called but not how many.

**(F)** Odds ratio for platelet protein detection (Burkhart et al.) per twofold platelet enrichment beyond megakaryocyte expression, mRNA half-life and nuclear:cytoplasmic ratio: 1.19 in Siphon-seq, 1.26 in the high-purity halves, 1.22 before purification, 1.27 and 1.28 in the low-purity halves. The gene-level association with protein detection is similar in all six datasets.

**(G)** Spearman ρ across gene sets between core-membership observed/expected and protein observed/expected (Burkhart et al.): 0.51 in Siphon-seq, 0.55 in both high-purity halves, 0.52 before purification, but 0.33 and 0.18 in the low-purity halves. The set-level association falls with contamination. PaxDb gives the same pattern.

**(H)** Sets called in each dataset relative to Siphon-seq, which calls 305 enriched and 186 depleted on the common universe. Right of zero, enriched sets: kept, newly enriched, and no longer enriched. Left of zero, depleted sets, drawn the same way. Public halves use a consensus call, the same direction in at least 10 of 20 draws. Moderate contamination keeps roughly half the Siphon-seq calls and adds many new ones: 173 of 305 enriched kept with 145 gained in P13 high purity, 171 with 148 gained in P14 high purity, and 168 with 184 gained before purification, of which 24, 24 and 37 carry an immune or inflammatory name. Depleted sets kept are 118, 118 and 99. The low-purity halves keep almost nothing (10 and 3 enriched, 22 and 6 depleted).

**(I)** $\log_2$ observed/expected of recurrent programs, columns ordered by leukocyte-marker RNA; dots mark sets significant in that dataset. Top block, 12 programs enriched in at least three datasets and higher there than in Siphon-seq (Toll-like receptor cascades, response to cytokine, signaling by interleukins, response to interleukin-1, Dectin-1 signaling and others). Bottom block, 12 programs depleted in Siphon-seq that are no longer depleted in at least two of the moderately contaminated datasets (tRNA aminoacylation, DNA integrity checkpoint, chromosome segregation, spliceosome and others). Asterisk, immune or inflammatory name.

#### Supplemental Figure 8. Effect of purity on gene-set sorting

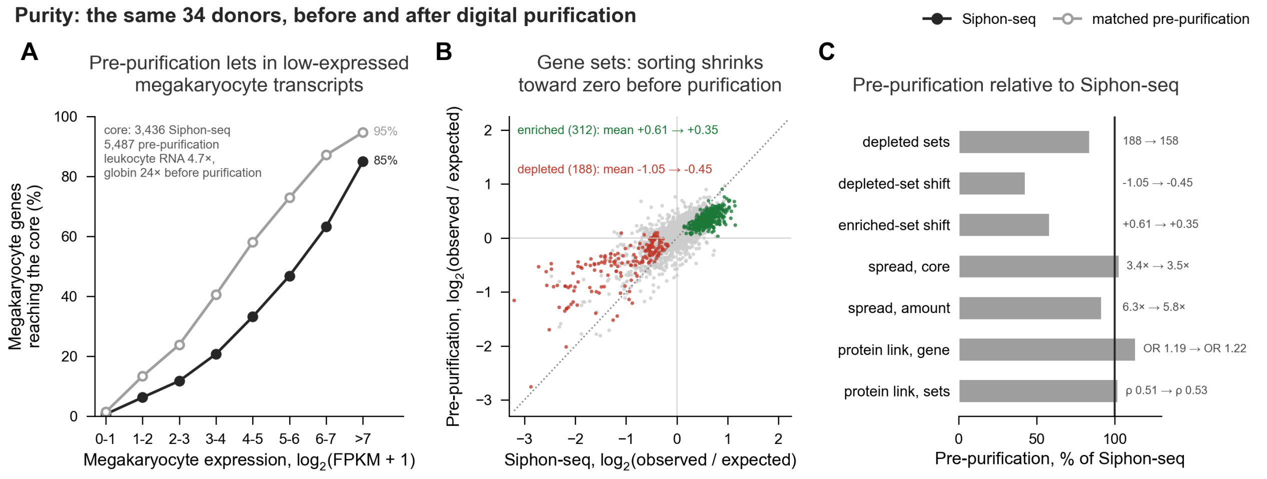


**Figure S8. Effect of purity on gene-set sorting.** The same 34 donors before and after removal of non-platelet partitions, so both arms share libraries and isolation protocol. Core membership in each arm uses the rule of the main text, mean CPM ≥ 2 and CPM > 2 in ≥ 90% of donors after TMM normalization.

**(A)** Percentage of megakaryocyte-quantified genes reaching the core, by megakaryocyte expression (BLUEPRINT, $\log_2$ FPKM + 1). Before purification roughly twice as many genes are admitted at every expression level, for example 40.6% against 20.7% at expression 3 to 4. The core holds 3,436 genes after Siphon-seq and 5,487 before it, while the pre-purification libraries carry 4.7 times the leukocyte-marker RNA (717 against 153 CPM) and 24 times the globin RNA (5,966 against 245 CPM).

**(B)** Core-membership enrichment of each of the 2,206 gene sets, Siphon-seq against pre-purification, both as $\log_2$(observed/expected) and clipped for display. Color, the Siphon-seq result (enriched, depleted, or not significant at Benjamini-Hochberg $q < 0.05$); dotted line, identity. Both signals shrink toward zero without purification: the 312 enriched sets move from a mean of +0.61 to +0.35 and the 188 depleted sets from −1.05 to −0.45.

**(C)** Pre-purification values as a percentage of Siphon-seq values for seven readouts, with the raw pair printed beside each bar. Set counts and mean shifts are simple ratios; the two spread measures use the excess over the null (variance ratio minus 1); the protein odds ratio is compared on the log-odds scale, so those three bars are not directly comparable with the others. Depleted sets 188 to 158, depleted-set shift −1.05 to −0.45, enriched-set shift +0.61 to +0.35, spread against the null for core membership 3.4-fold to 3.5-fold and for transcript amount 6.3-fold to 5.8-fold, protein odds ratio 1.19 to 1.22, and set-level ρ between sorting and protein 0.51 to 0.53. Purification changes which sets are sorted and the size of the shift, while the association with platelet protein is unchanged.

### Supplemental Figure 9. In silico titration

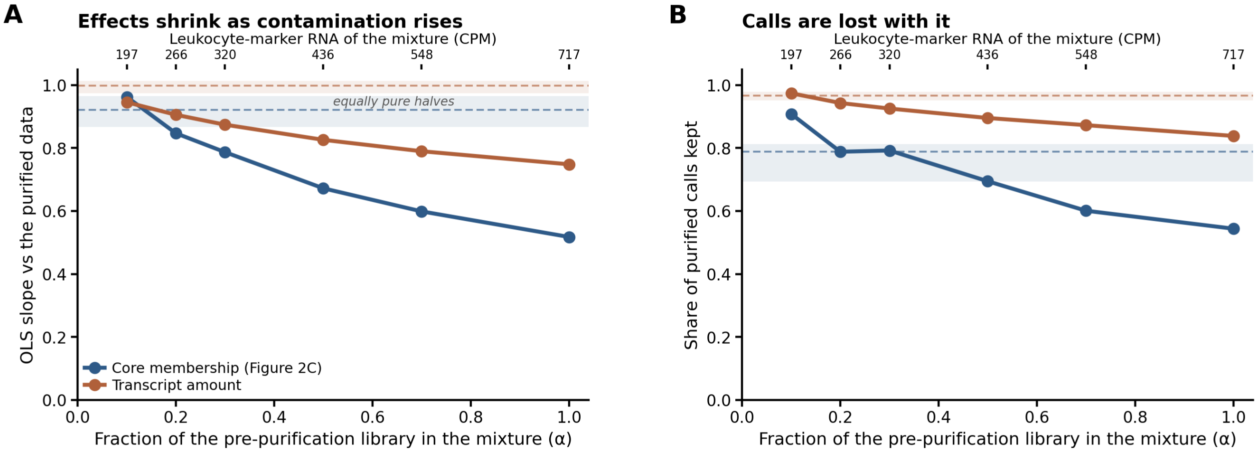


**Contamination alone reproduces the loss of the gene-set sorting results.** The same 34 donors were mixed in TMM-CPM space as α × their pre-purification library + (1 − α) × their Siphon-seq library, and each mixture was scored against α = 0. Donors, platform, library chemistry and processing are identical across the series, so contamination is the only variable. The top axis gives the leukocyte-marker RNA of each mixture, for comparison with datasets of similar purity. Two readouts are shown throughout: core membership (the set-wise enrichment of core genes) and transcript amount.

**(A)** Attenuation. OLS slope of each mixture's effects against the same effects in the purified data; 1.0 means effects are recovered at full size. Core membership falls from 0.96 at α = 0.1 to 0.52 at α = 1, and transcript amount from 0.95 to 0.75. **(B)** Loss of calls. Share of the significant calls made in the purified data that the mixture still makes. Core membership falls from 0.91 to 0.54, transcript amount from 0.97 to 0.84.

In both panels, the dashed lines and shaded bands are the noise floor: the median and interquartile range of the same statistic computed between two halves of equally pure data (20 batch-balanced 17 + 17 splits of the purified donors). Two equally pure halves already give a slope below 1, because noise in the x-axis dataset flattens an OLS fit; attenuation within the band is therefore attributable to measurement noise. In (A), core membership leaves that band between α = 0.1 and α = 0.2, at roughly 200–270 CPM of leukocyte-marker RNA, while transcript amount, whose band is narrower, is already below it at α = 0.1 (slope 0.95 against a lower quartile of 0.97) but declines more slowly thereafter (0.75 at α = 1 against 0.52). In (B), transcript amount leaves its band between α = 0.1 and α = 0.2, whereas core membership, whose split-half band is wide, leaves it only between α = 0.5 and α = 0.7.

α = 1 reproduces the pre-purification libraries exactly (717 CPM leukocyte-marker RNA, slope 0.517).

### Supplemental Figure 10. Independent sepsis cohort (GSE252275)

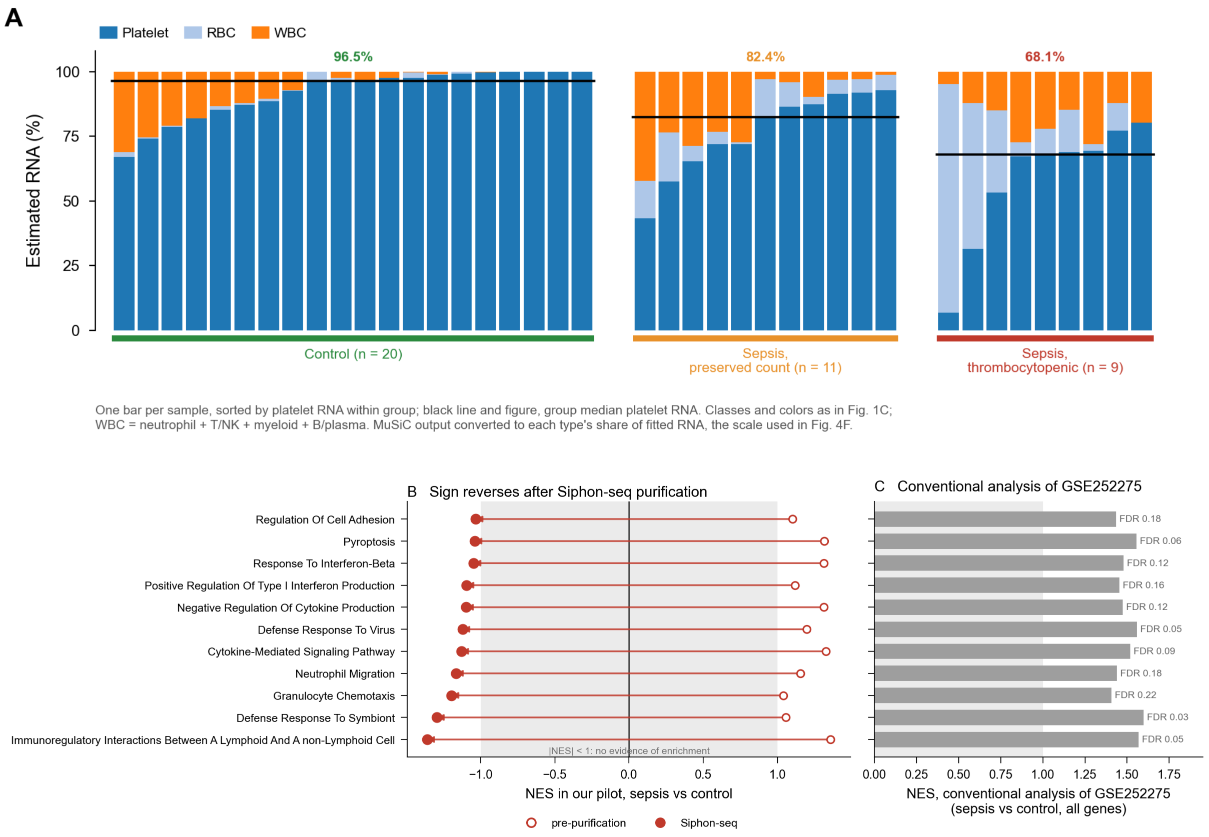


**Figure S10. Contamination in an independent sepsis cohort (GSE252275) increases with illness severity and overlaps pathways reported as enriched in sepsis.** GSE252275 is a published platelet RNA-sequencing cohort of 40 samples: 20 controls, 11 patients with sepsis and a preserved platelet count, and 9 patients with sepsis and thrombocytopenia.

**(A)** Estimated RNA composition of every sample, one bar per sample, sorted by platelet RNA within group, with leukocyte the sum of the neutrophil, T/NK, myeloid and B/plasma estimates. Composition is MuSiC output converted to each cell type's share of fitted RNA, rather than the raw cell proportion. Line and figure above each group, the group median platelet RNA: 96.5% in controls, 82.4% in sepsis with a preserved platelet count and 68.1% in sepsis with thrombocytopenia, with median leukocyte RNA rising from 2.4% to 9.6% to 14.9% and red-cell RNA from 0.4% to 5.9% to 10.7%. The spread widens with severity, from 67.1 to 100% platelet RNA in controls to 6.9 to 80.3% in the thrombocytopenic group. Contamination therefore differs systematically between the groups being compared. No statistical test is drawn in this panel.

**(B)** Pathways whose direction reverses once contaminating RNA is removed. Of the pathways scoreable in both arms of our paired sepsis pilot, 354 are positively enriched for sepsis against control before purification and negatively enriched after; 29 of these are also reported as enriched at FDR < 0.25 by a conventional analysis of GSE252275. Of those 29, the 11 drawn here reach |NES| ≥ 1 on both sides; the shaded band on each axis marks |NES| < 1. Open circle, pre-purification; filled circle, Siphon-seq; arrow, the shift between them, drawn for our pilot and not for GSE252275. Pilot enrichment scores are from preranked GSEA against GO Biological Process 2023 and Reactome 2022 with 10,000 permutations, the same runs as the sepsis pilot analysis (4 patients with sepsis against 4 controls before purification, 3 against 3 after), and 2,485 terms are scoreable in both arms. All 11 are immune or inflammatory terms, led by immunoregulatory lymphoid interactions (+1.36 to −1.36), defense response to symbiont (+1.05 to −1.29), granulocyte chemotaxis (+1.04 to −1.20) and neutrophil migration (+1.16 to −1.16); their leading-edge genes are a median of 16.3% myeloid-specific.

**(C)** The same 11 pathways as reported by a conventional analysis of the deposited GSE252275 counts, with no contamination removal: NES for sepsis against control over all genes, with the false-discovery rate beside each bar. All 11 are reported as enriched, with NES 1.41 to 1.60 and FDR 0.03 to 0.22, only two of them below 0.05 (0.035 and 0.046; the next, 0.052, is printed as 0.05).

These are our preranked analyses of the deposited counts (edgeR quasi-likelihood differential expression, genes ranked by $\log_2$ fold change × $-\log_{10}$ P, the same two libraries, 1,000 permutations), not values from the source publication.

#### Supplemental Data files

The following are uploaded separately as Excel workbooks.

**Supplemental Data 1.** Per-sample MuSiC estimates for the 3,689 public samples.

**Supplemental Data 2.** Per-donor partition calls: % GEMs and % RNA by class, RNA/GEM ratios, MuSiC platelet % before and after purification, both cohorts.

**Supplemental Data 3.** Core platelet transcriptome: all 38,606 genes with mean CPM, % donors detected, core / contextual / below-detection call, chromosome, Y-linked flag, megakaryocyte expression.

**Supplemental Data 4.** Gene-set core membership, observed versus expected.

**Supplemental Data 5.** Gene-set changes with purification.

**Supplemental Data 6.** Protein detection by core membership.

**Supplemental Data 7.** Antithrombotic drug-target pairs (full ChEMBL mapping).

**Supplemental Data 8.** Sepsis pilot pathway results.

**Supplemental Data 9.** GSE252275 results.

**Supplemental Data 10.** Comparison with earlier platelet gene sets.

**Supplemental Data 11.** Pre-purification versus Siphon-seq gene-level comparison: all 4,558 tested genes, and the 20 most increased in each direction with their lineage categories.
